# Displacement of extracellular chloride by immobile anionic constituents of the brain’s extracellular matrix

**DOI:** 10.1101/2022.12.28.522113

**Authors:** Kieran P Normoyle, Kyle P Lillis, Kiyoshi Egawa, Melanie A McNally, Mousumi Paulchakrabarti, Biswa P Coudhury, Lauren Lau, Fu Hung Shiu, Kevin J Staley

## Abstract

GABA is the primary inhibitory neurotransmitter. Membrane currents evoked by GABA_A_ receptor activation have uniquely small driving forces: their reversal potential (E_GABA_) is very close to the resting membrane potential. As a consequence, GABA_A_ currents can flow in either direction, depending on both the membrane potential and the local intra and extracellular concentrations of the primary permeant ion, chloride (Cl). Local cytoplasmic Cl concentrations vary widely due to displacement of mobile Cl ions by relatively immobile anions. Here we use new reporters of extracellular chloride (Cl^-^_o_) to demonstrate that Cl is displaced in the extracellular space by high and spatially heterogenous concentrations of immobile anions including sulfated glycosaminoglycans (sGAGs). Cl^-^_o_ varies widely, and the mean Cl^-^_o_ is only half the canonical concentration, i.e. the Cl concentration in the cerebrospinal fluid. These unexpectedly low and heterogenous Cl^-^_o_ domains provide a mechanism to link the varied but highly stable distribution of sGAGs and other immobile anions in the brain’s extracellular space to neuronal signal processing via the effects on the amplitude and direction of GABA_A_ transmembrane Cl currents.

**Key Points Summary:**

- Extracellular chloride concentrations in the brain were measured using a new chloride-sensitive organic fluorophore and two photon fluorescence lifetime imaging.
- *In vivo*, the extracellular chloride concentration was spatially heterogenous and only half of the CSF chloride concentration
- Stable displacement of extracellular chloride by immobile extracellular anions was responsible for the low extracellular chloride concentration
- The changes in extracellular chloride were of sufficient magnitude to alter the conductance and reversal potential of GABA_A_ chloride currents
- The stability of the extracellular matrix, the impact of the component immobile anions including sulfated glycosaminoglycans on extracellular chloride concentrations, and the consequent effect on GABA_A_ signaling suggests a previously unappreciated mechanism for modulating GABA_A_ signaling.

## Introduction

GABA_A_ synapses comprise the primary output for inhibitory interneurons in the brain (Hu *et al*., 2014; Tremblay *et al*., 2016). A unique feature of GABA_A_ synapses are the very small driving forces for ion flux through open GABA_A_ receptor-operated membrane channels. The reversal potential for the primary permeant ion, chloride, is usually within a few mV of the neuronal resting membrane potential (RMP) (Thompson *et al*., 1988) and may even be positive to RMP (Staley & Mody, 1992; Tyzio *et al*., 2008). This low driving force makes possible a novel form of synaptic plasticity at GABA_A_ synapses: the direction of current flow, i.e. the effect of GABA_A_ receptor activation on the resting membrane potential, can be reversed by low millimolar alterations in the local chloride concentration (Coombs *et al*., 1955). Such physiological variance is routinely observed in individual studies (Ebihara *et al*., 1995; Huberfeld *et al*., 2007) and is likely to underlie the 25 mV range of reported group mean reversal potentials in 20 recent studies of evoked GABA_A_ currents recorded with gramicidin perforated patch electrodes in area CA1 of the hippocampus (tabulated in (Rahmati *et al*., 2021). This variance has substantial effects on synaptic signaling: the reversal potential of GABA_A_ currents evoked in a single pyramidal cell by stimulation of 8 different presynaptic interneurons also varied by over 20 mV (Rahmati *et al*., 2021).

We have proposed that the intracellular chloride concentration (Cl^-^_i_) is not monotonic across the neuronal cytoplasm, but rather varies inversely with the local concentration of macromolecular anions that comprise the vast majority of intracellular anions (Glykys *et al*., 2014a; Rahmati *et al*., 2021). This phenomenon is known as Donnan exclusion (Marinsky, 1985; Helfferich, 1995; Fatin-Rouge *et al*., 2003; Epsztein *et al*., 2018; Gao *et al*., 2022). Mobile ions, in this case Cl, are repelled by relatively immobile ions such as proteins with negative surface charges. Many proteins such as gephyrin that are associated with GABA_A_ synapses bear large negative surface charges (Sola *et al*., 2001), suggesting that variance in the GABA_A_ reversal potential (E_GABA_) is determined by local fixed cytoplasmic charge densities. However, E_GABA_ is not determined solely by Cl^-^_i_ but rather by the ratio of intra- to extracellular permeant ions. This raises the possibility that spatial variance in the extracellular Cl concentration (Cl^-^_o_) could also be an important determinant of E_GABA_.

The displacement of Cl^-^ by fixed negative charges should also occur in the extracellular space, where the high negative charge density of several classes of functionalized macromolecules are anchored to the cell membrane either directly or indirectly. Gangliosides typically bearing one to three – or more, though rarely – carboxylate groups in the form of sialic acid groups are present in the extracellular space of human cerebral cortex (Schnaar *et al*., 2014) and are concentrated within the synaptosomal fraction (Svennerholm, 1980). Together with the polysulfated glycosaminoglycans (GAGs) that are a major constituent of the extracellular matrix (Karamanos *et al*., 2021), which have been reported to be as high as the Cl^-^ concentration in the CSF (Urban *et al*., 1979; Chahine *et al*., 2005; Morawski *et al*., 2015), we calculate that total extracellular fixed anions are in excess of 50mM (see Methods; Schnaar *et al*., 2014). Thus some to all of the local Cl^-^_o_ could be displaced, depending on local anionic density and the degree of GAG sulfation. Such displacement has been most clearly demonstrated in the chondroitin sulfate-rich matrix of cartilaginous tissue, where the negative charge density of the extracellular matrix can reach 400 meq / liter (Lesperance *et al*., 1992), but dozens of mM of Cl^-^_o_ are also displaced by sulfated GAGs in tissues such as skin (Titze *et al*., 2004). Mucous, which comprises the extracellular space for airway epithelial cells, is also rich in polysulfated GAGs that displace about half of the Cl^-^_o_ (Reuter *et al*., 1998). Given this evidence for sulfated GAGs displacing Cl^-^_o_ and the commercial availability of exogenous matrix metalloprotease (MMP) analog chondroitinase, we chose to focus on the role of extracellular sulfated GAGs and, in particular, the role of chondroitin sulfate.

Many findings suggest that the brain Cl^-^_o_ is lower than the canonical value. Xray emission spectroscopy of the extracellular space has demonstrated sulfate concentrations as high as 100 mM (Morawski *et al*., 2015). Astrocytic buffering of Cl^-^_o_ (Egawa *et al*., 2013) would not be necessary if Cl^-^_o_ were indeed > 100 mM. The gating of Group II and III metabotropic glutamate receptors, kainate receptors, acid-sensing membrane ion channels, and the transport rate of the GlyT1 glycine transporter are all strongly modulated by Cl^-^_o_ over the entire range of 1 - 100 mM (Kuang & Hampson, 2006; Plested & Mayer, 2007; Kusama *et al*., 2010; DiRaddo *et al*., 2015; Tora *et al*., 2018; Zhang *et al*., 2021). Such strong modulation by low Cl^-^_o_ would not make sense if Cl^-^_o_ was uniformly > 100 mM.

Studies of Cl^-^_o_ are few, particularly in live mammalian tissue, and have been carried out with ion-sensitive electrodes that sample only one microenvironment (Nicholson & Kraig, 1975). Our prior work suggested that the local chloride concentration varies widely as a function of the local concentration of immobile anions (Rahmati *et al*., 2021). Earlier results from our group (Glykys *et al*., 2014a) have suggested that physiological Cl^-^_o_ is lower than that measured with ion-sensitive electrodes, and we hypothesized that the large sample sizes provided by fluorometric imaging techniques might provide the most robust measure of the extracellular chloride distribution.

Although a fluorometric study of Cl^-^_o_ would provide a noninvasive means to establish the distribution of chloride in the extracellular space, no suitable Cl-sensitive organic fluorophores exist, and transgenic fluorophores such as SuperClomeleon (Grimley *et al*., 2013) and CloPhensor (Sato *et al*., 2017) are not distributed in the extracellular space. We therefore developed a chloride-sensitive fluorophore (Normoyle *et al*., 2024) from previously described organic backbones (Biwersi *et al*., 1992; Biwersi *et al*., 1994) that satisfied the following criteria.

First, the fluorophore should be sensitive to Cl in the range of the extracellular chloride concentration i.e., up to 100 mM; available fluorophores have been optimized for the low millimolar chloride concentrations of the cytoplasm. Second, the fluorophore should report Cl independently of the concentration of fluorophore. Ratiometric fluorophores suitable for the extracellular space do not exist, so we used **<u>F</u>**luorescence **<u>L</u>**ifetime **<u>Im</u>**aging (FLIM) (Kaneko *et al*., 2004) to remove the dependence of the fluorescence emission on fluorophore concentration. Third, the fluorophore should be constrained to the extracellular space, for which dextran conjugation is optimal (Xiao *et al*., 2008; Tønnesen *et al*., 2018). Fourth, the fluorophore should be excited by wavelengths that are not damaging to brain tissue, i.e. below the ultraviolet range and in the visible or infrared range. Fifth, many Cl sensors are also sensitive to pH (Kuner & Augustine, 2000; Grimley *et al*., 2013), so a pH-insensitive fluorophore would be ideal. Finally, the fluorophore needs to be sufficiently bright so that its emission can be reliably distinguished from autofluorescence.

Here, we utilize a newly synthesized Cl-sensitive fluorophore that meets these 6 criteria to demonstrate: stable Donnan exclusion of chloride by immobile extracellular anions such as sulfates fixed to GAG-like carbohydrate polymers; directly measured Cl^-^ of *in vitro* and *in vivo* brain tissue that is much lower than cerebrospinal or plasma chloride; a remarkable spatial variance in Cl^-^; and, as evidence that Cl^-^ is determined by Donnan exclusion of Cl by sulfated moieties such as GAGs, Cl^-^ increases after release of immobile sulfates by exogenous chondroitinase.

## Methods

### Ethical Approval

All animals are housed in a facility with on-call veterinary team and have free access to food water in accordance with MGH IACUC animal welfare board regulations, conforming to Grundy 2015 principles and regulations (protocol 2018N000221). Isofluorane anaesthesia was induced by inhalation through a nosecone at 5mL/min and maintained at 1- 2mL/min for the duration of the experiment, checking frequently for signs of adequate anaesthesia. All surgeries performed were acute (non-survival) experiments after which mice were administered maximum anaesthesia (5mL/min isofluorane) and euthanized by decapitation.

### General

Chemicals were purchased from Sigma-Aldrich (St Louis, MO) unless otherwise noted. Solutions are brought up in distilled water unless otherwise noted. Standard aCSF was brought up in 18MΩ water and consisted of (in mM): NaCl 126; KCl 3.5; CaCl_2_ 2; MgCl_2_ 1.3; NaH_2_PO4 1.2; Glucose 11; NaHCO_3_ 15. Low chloride aCSF variant was identical except that sodium gluconate was substituted for NaCl, resulting in 10mM Cl^-^. Calibration solutions were made by mixing appropriate proportions of standard aCSF with low chloride aCSF such that a chloride and gluconate sum total of 136mM was maintained while Cl^-^ ranged from 10 to 136mM. Colorimetric assays were read using a Wallac Victor-2 1420 spectrophotometer with a halogen continuous wave light source and spectral line filters at listed wavelength +/- 5 to 10nm. Centrifugation steps were accomplished in a tabletop Eppendorf 5417R microcentrifuge. Reflux apparatus for synthesis consisted of a recirculating cooling bath (Fisher Scientific) filled with ethylene glycol cooling a 24/40 double-lumen coiled reflux tube (Ace Glassware, Vineland, NJ) with a 500mL round bottom flask (Corning) heated with a heating mantle regulated by a timed power controller (Glas-Col #O406 and #104A PL312, respectively).

### Synthesis of ABP-dextran

The process of optimizing a bright, redshifted, pH-insensitive fluorophore that is responsive to chloride concentration over a dynamic range inclusive of expected extracellular chloride concentrations is beyond the scope of this paper and has been published separately (Normoyle *et al*., 2024). Synthesis techniques closely followed those of the Verkman group when synthesizing related compounds (Verkman, 1990; Biwersi *et al*., 1992) and conjugating to dextran (Biwersi *et al*., 1992). Briefly, N(4-***A***mino***B***utyl)***P***henantridinium (ABP) was synthesized by a reflux of equimolar amounts of phenanthridine and N(4- bromobutyl)phthalimide in acetonitrile followed by a second reflux in 6N HCl to hydrolyze phthalate. Product was twice recrystallized from 95% ethanol before conjugation to cyanogen bromide-activated 10,000g/moL (10KDa) dextran, producing 10KDa ABP-dextran with an approximate molar labeling ratio of 4.4.

### Sulfation of agarose

We modified the method of Fuse and Suzuki (Fuse & Toshiya, 1975) to add sulfate groups to the repeating disaccharide units of agarose. The goal of maximizing sulfation must be balanced with the consequent erosion of gel properties such as gelation and water holding capacity; in our hands, the maximum sulfation we could achieve while retaining gelation at 5% (w/v) was when using a 6:1 molar ratio of sulfuric acid to agarose disaccharide. When cast at 5% (w/v), this worked out to ≈50mM sulfate in resultant gels (49.2+/-0.289mM; for sulfate measurement see next section). This roughly matches calculated estimates of fixed negative charges from a collection of bulk biochemical studies quantifying the content of sulfate groups (proteoglycans and sulfatides) and carboxylic acid groups (sialic acid-bearing glycolipids and proteins) in brain (Schnaar *et al*., 2014). Schnaar et al. (2014) summarized the components of the brain’s extracellular matrix as follows: proteoglycans represent 0.45mM disaccharide monovalent negative charge equivalents and sulfatides represent 5.8mM charge equivalents, while sialic acid groups on gangliosides account for 3.9mM, and sialic acid bound to protein comprises 1.3mM. In total, these distinct sources of extracellular negative charge bound to lipid, protein, and sugar components of the extracellular matrix amount to 11.45mM sequestered into the 20% of total volume that is extracellular. This gives a total expected fixed extracellular anionic charge equivalent of 57.3mM in the brain’s extracellular space. Proteoglycans, including chondroitin sulfate, account for 2.3mM of total extracellular fixed charge (0.45mM corrected for extracellular space being 20% of total). Despite this, we focus on chondroitin sulfate digestion of live tissue due to experimental accessibility and the ability to use commercially available chondroitinase. It is also for practical reasons that we model the complex extracellular matrix using sulfated agarose because of the experimental expediency of adding sulfate to a polymerizable sugar. We stress again that there is a limit to how much sulfate can be added to agarose while retaining its ability to polymerize, and the following describes a sulfated agarose with sulfate content near or at this limit. Briefly, agarose was added to a well-dried Erlenmeyer flask and dissolved in DMSO at a concentration of 4%(w/v) at room temperature before being cooled to 4°C with stirring. Addition of 4:1 mixture of acetic anhydride / glacial acetic acid, also pre-chilled to 4°C, was added to agarose/DMSO mixture (0.4 volumes) in 4°C coldroom. Vigorous mixing by handheld rotation was necessary to thin the gelatinous mixture to a slurry and allow stirring at 4°C for 15 minutes; gentle heat transfer from experimenter’s hands helped loosen the initial gel but care is taken to return the mixture to 4°C. Sulfuric acid (15.6M) is then added slowly at a 6:1 molar ratio to agarose repeating disaccharide units, taking care to keep temperature between 4-10°C. Mixture is then moved to ambient temperature and allowed to stir for 15 minutes before reaction is stopped by neutralization with 10M NaOH after again cooling to 4°C and maintaining temperature between 4-10°C in coldroom. Sulfated agarose is then dialyzed at ambient temperature in 12-14KDa dialysis tubing (SpectraPor S432703) against 15 volumes of 50mM Tris pH 7.4 once, then against distilled water changed twice daily until dialysate clarifies. Agarose is then precipitated in 95% ethanol cooled to 4°C >4hours; over-dialysis may hinder ethanol precipitation due to a lack of sufficient counter-ion presence. Precipitate is then vacuum filtered and oven-dried at 50°C. Material is loosened with polished glass pestle and stored at ambient temperature in sealed glass vial until use. Note that careful temperature and pH control is necessary throughout exothermic processes to maintain gelation characteristics of the product.

### Measurement of sulfate concentration in polysaccharide gels

BaCl_2_-gelatin turbidimetric method (Torres *et al*., 2021) was used to quantify the sulfate content of gel materials. Sulfated agarose product or commercially available agarose or agar powder was dissolved in 0.5M HCl at a concentration of 5% (w/v) in 2mL screw-top microtubes with gasket (Fisher Scientific). Tubes are then placed in a heat block at 100-110°C for 3hours with periodic vortex mixing, hydrolyzing the sugar backbone and releasing sulfate. All samples had varying amounts of insoluble dark brown precipitated sugar, which was clarified at 20,000 x g for 10 minutes. Standards are diluted from 1M Na_2_SO_4_ stock in 0.5M HCl. BaCl_2_-gelatin reagent is freshly prepared by dissolving 30mg gelatin in 10mL distilled water at 80°C for 10minutes, then promptly adding 100mg BaCl_2_ and vortex mixing. BaCl_2_-gelatin is then allowed to cool passively before use. Samples are prepared by combining 1 part sample (diluted as desired), 1 part BaCl_2_-gelatin, and 3 parts 0.5M HCl. Assay is performed in a glass-bottom 96-well plate reading absorbance at 405nm. Multiple sample dilutions were prepared and concentrations were calculated only from dilutions whose readings were well within the dynamic range of the standards.

### Measurement of Cl^-^ in agar or agarose gels (chemical method)

Chloride was measured colorimetrically using mercury(II) thiocyanate (Hg(SCN)_2_) / iron(III) nitrate (Fe(HNO_3_)_3_) absorbance method of Florence and Farrar (Florence & Farrar, 1971) to infer Cl^-^ from Fe(SCN)_3_ production through the two-step reaction,

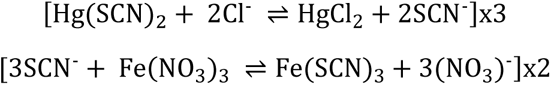

reading Fe(SCN)_3_ absorbance at 490nm. Briefly, agar or agarose gels cast at 5% (w/v) in 3.5cm Petri dishes were drained by gravity and then Kimwipes were used to gently wick away remaining excess fluid at sides of dish. Similar amounts of each sample are transferred to 2mL screw-top microtubes with gasket (Fisher Scientific) and 1 gel volume, estimated by weight, of 7.8M nitric acid (HNO_3_) is added (final [HNO_3_] = 3.9M). Samples are then placed in a heat block at 100-110°C for 3hours with periodic vortex mixing, hydrolyzing the sugar backbone to break down the gel lattice. All samples had varying amounts of insoluble dark brown precipitated sugar, which was clarified at 20,000 x g for 10 minutes. A portion of supernatant or KCl standard, diluted with 0.3M HNO_3_, was combined with one-tenth volumes of 0.4M Fe(NO_3_)_3_ (in 4M nitric acid) and saturated Hg(SCN)_2_ (in ethanol), triturated, and allowed 20minutes for reaction to reach endpoint. Stock solutions are used within 1week. Multiple sample dilutions were prepared and concentrations were calculated only from dilutions whose readings were well within the dynamic range of the standards. Final concentrations are adjusted to account for dilution from original gel sample volume.

### Fluorescence Lifetime Imaging (FLIM)

Time-correlated single-photon counting (TCSPC) FLIM measurements were obtained using a custom laser-scanning two-photon rig driven by custom ScanImage software (MBF Bioscience, Williston, VT) equipped with a MaiTai Ti:Sapph laser (SpectraPhysics) and high-sensitivity PMT detectors (Hamamatsu C6438-01) as described previously (Normoyle *et al*., 2024). Briefly, ABP-dextran was excited at 760nm and emitted photons were subjected to a 445/58 bandpass filter (Chroma) prior to detection using a high-sensitivity PMT. ABP-dextran was used at 500microgram/mL (50μM) in aCSF unless otherwise noted. A fluorescence lifetime is calculated for each pixel by measuring the time it takes for an emitted photon to be detected, binning these photons according to arrival time, and to these data a first-order exponential equation is fit to derive the rate constant representing the fluorescence lifetime (Gehlen, 2020). The presence of a quencher will shorten fluorescence lifetime to an extent which is linearly related to quencher concentration through the Stern-Volmer equation,

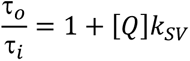

where *τ_o_ / τ_i_* represents the unquenched lifetime divided by the shortened lifetime at a given quencher concentration ([*Q*]) (Gehlen, 2020). The Stern-Volmer constant (*k_SV_*) of the resultant line relates the concentration of quencher to the shortening of fluorescence lifetime. Photons were collected over an acquisition period of 60 seconds (*in vivo* and agarose gels) to 90 seconds (*in vitro* slice cultures) and the lag between incident laser pulse and resulting photon detection recorded. The population of photons in each individual pixel is fit to an exponential to yield a lifetime value and summed to yield an intensity value using the FLIMJ plug-in for ImageJ. Further image processing, calculations, and statistics were accomplished using built-in functions and custom routines developed in Matlab (v2018b and v2022a; Mathworks, Natick, MA). For composite image generation, see *Image processing in Matlab* section below.

### Calibration of ABP-dextran

ABP-dextran is synthesized, conjugated, and calibrated as described previously (Normoyle *et al*., 2024). Briefly, we calibrated 50μM ABP-dextran against aCSF with varying chloride concentrations by preparing standard aCSF (in mM: NaCl 126; KCl 3.5; CaCl_2_ 2; MgCl_2_ 1.3; NaH_2_PO4 1.2; Glucose 11; NaHCO_3_ 15) and a low-chloride aCSF in which NaCl was replaced with sodium gluconate. We mixed appropriate amounts of each to vary chloride concentrations (10-136mM) while maintaining the sum of sodium chloride and sodium gluconate at 136mM. Although *k_SV_* values did not differ appreciably between ABP- dextran batches nor when measured using different FLIM-capable microscopes, out of an abundance of caution we used a single batch of ABP-dextran and obtained FLIM calibration parameters specific for this batch of fluorophore and the custom microscope used for all experiments reported in this manuscript. Because calibrations done in aCSF have a minimum Cl^-^ of 10mM as described, the unquenched lifetime (*τ_o_*) used for Stern-Volmer calculations is approximated as the y-intercept (Cl^-^ = 0mM) of a best-fit curve of the raw calibration data; empirically, a biexponential fit was used. Each concentration of Cl^-^ in the calibration curve has a normally distributed set of ABP-dextran lifetime values with a near-constant relative standard deviation (RSD = standard deviation divided by mean) (Normoyle *et al*., 2024). All calculations are otherwise done as described under *Fluorescence Lifetime Imaging (FLIM)* section above.

### Animals

All animal protocols were approved by the Massachusetts General Hospital Institutional Animal Care and Use Committee (protocol 2018N000221). Wild type mice (C57bl/6; Jackson Labs 000664) of either sex were used for this study. Mouse pups remained in the home cage with the dam under standard husbandry conditions until postnatal day 6 to 8 (P6–8) when organotypic slice cultures were prepared, or until P26-30 when a cortical window was surgically placed.

### In vitro organotypic slice cultures and imaging

Organotypic hippocampal slice cultures were prepared either as glass-mounted (Berdichevsky *et al*., 2016) or membrane insert-mounted (Stoppini *et al*., 1991) cultures. Briefly, in either case hippocampi are obtained from P6-8 mice and cut to 400μm thick slices. These are then gently transferred to a 6-well dish containing either a membrane insert (Millipore) or a poly-L-lysine coated coverslip (Electron Microscopy Sciences), are fed twice weekly with 1mL neurobasal-A media supplemented with 500μM Glutamax, 2% B-27, and 0.03mg/mL gentamycin (all from Invitrogen), and are incubated at 35°C in a 5% CO_2_ incubator. Cultures are typically imaged between DIV 12-21 in aCSF warmed to 33°C and bubbled with 95%O_2_ / 5% CO_2_ perfused at a rate of approximately 100mL/hour unless otherwise noted. Slices were pretreated with media containing 500μg/mL ABP-dextran in incubator for 1-2 hours prior to perfusing with aCSF containing 136mM chloride and the same concentration of ABP-dextran, plus 100nM tetrodotoxin (TTX). Slices were allowed to equilibrate in perfusate for 20-30 minutes before imaging, equaling at least 2 hours of total exposure to ABP-dextran prior to initiation of imaging. Two photon images were captured on a perfusion-enabled stage with a 20x water-immersion objective, NA 0.95, and analyzed as described (see *Image processing in Matlab* section below).

### *In vivo* murine cortical window imaging

Using a modified method based on that of Che and De Marco Garcia (Che & De Marco García, 2021), we placed cortical windows in young adult mice (P26-30) for acute (non-survival) imaging in accordance with Massachusetts General Hospital Institutional Animal Care and Use Committee policies and procedures (protocol 2018N000221). Briefly, anaesthetized mice were immobilized with standard ear bars and nosepiece until custom headbar is placed. A section of scalp is removed to expose a roughly 1cm section of skull. Acrylic dental cement powder (Lang Dental, Wheeling IL) mixed with cyanoacrylate adhesive is used to fix the headbar to the exposed skull at 4 o’clock to Bregma, creating a 5mm diameter working area just lateral to midline. A 2.5mm round section of skull and underlying dura is removed and a 3mm No. 1 coverslip is placed over the exposed cortex. Coverslip has 10μL concentrated ABP-dextran / agarose mixture (1%w/v low gelation temperature agarose, Sigma-Aldrich, with 10mg/mL ABP-dextran in aCSF) pipetted from 42°C heat block immediately before inversion and placement over cortex. One hour is allowed before image acquisition for ABP- dextran diffusion into cortex from overlying agarose (between cortex and coverslip), producing signal similar to that seen with 50μM ABP-dextran in vitro. During this time the warmed breadboard with immobilization apparatus and anesthetized mouse is transferred to the microscope table for *in vivo* imaging. This custom-made gantry-type microscope accommodates the breadboard as a stage for *in vivo* imaging. Two photon images were captured with a 16x water-immersion long focal distance objective, NA 0.80, and were analyzed as described (see *Image processing in Matlab* section below).

### Agarose or agar gel Cl^-^ determination using ABP-dextran FLIM

Cl^-^ within an agar or agarose gel was measured using the two-photon rig described above in the *in vivo* section, except with a 4x (NA 0.28) objective. Using an 8-well microchamber with silicone dividing gasket (Ibidi, Grafelfing, Germany), gels were cast in aCSF containing ABP-dextran and subsequently transferred to a single Petri dish where they were equilibrated with aCSF containing ABP- dextran for at least 1 hour prior to imaging. FLIM measurements were taken as described, generally one field per quadrant at a depth of between 50-100microns. Importantly, the gasket separating gels of differing compositions was removed such that all gels in an experiment were bathed in the same aCSF with the same Cl^-^. Where gels were kept for more than 1 day, they were prepared in aCSF without glucose and kept refrigerated between readings to prevent fouling.

### Chondroitinase treatment of organotypic hippocampal slice cultures

Chondroitinase ABC (ChABC; AMSBIO, #E1028-02) was brought up in aCSF containing 0.1% bovine serum albumin (BSA) to a concentration of 1U/mL. Slices grown on membrane inserts were treated for 1hour either with aCSF containing ChABC or sham aCSF containing only the 0.1% BSA vehicle. Both experiment and control had 0.5mg/mL ABP-dextran (50μM), and FLIM imaging was done using the two-photon rig described in the *in vivo* section above. At the conclusion of imaging, the aCSF was collected for detection of liberated chondroitin sulfate.

### Measurement of sulfated glycosamioglycans (sGAGs) liberated in aCSF

Blyscan 1,9- dimethylmethylene blue-based kit (Biocolor, #B1000) was utilized for measurement of sGAGs including chondroitin sulfate proteoglycans (CSPGs). After treatment with aCSF containing ChABC or BSA vehicle as detailed above, aCSF was collected and measured as directed against standards of bovine trachea chondroitin (Blyscan kit standard) with appropriate amounts of 10x stocks of aCSF and 0.1% BSA to obtain concentrations identical to samples. 1,9- Dimethylmethylene blue reagent absorbance is read at 650nm.

### Measurement of chondroitin sulfate by liquid chromatography / mass spectroscopy (LC-MS)

aCSF containing either chondroitinase or vehicle was recovered after incubation with slice (see above), volume recovered carefully noted before freezing at −20°C, and shipped over dry ice with blinded labels for analysis. The samples were filtered using 3K MWCO spin filter units (Pall Corporation Nanosep with 3K Omega Ref: OD003C34) at 14000 RPM at 10°C for 20minutes. The flow through containing CS disaccharides was dried in speed vac. The dried flow through were reconstituted in 200µl ultrapure distilled water (Invitrogen, Cat # 10977-015) and 100µl was dried taken for ^12^C_6_ aniline tagging by reductive amination method (Lawrence *et al*., 2008). Briefly, the CS disaccharides were dissolved in 17μl of ^12^C_6_ aniline followed by addition of equal volume of freshly prepared 1 M NaCNBH_3_ (Sigma-Aldrich) in dimethyl sulfoxide: acetic acid mixture (65:35, v/v). The reaction mixture was kept at 70°C for 45min and then transferred to dry oven at 37°C for overnight reaction (16 h). After aniline tagging samples were dried in speed-vac for 48h to remove excess reagents. The samples were then dissolved in 17µL of LC-MS grade water (Honeywell), centrifuged at 7000g for 5min. 5µL of the sample was taken and mixed with 5µL reconstitution buffer containing known amount of internal standard (5µl of the reconstitution buffer contains, 1 μl of 10X LC-MS buffer-A (80 mM acetic acid and 50 mM dibutylamine; Sigma-Aldrich), 2 μl of ^13^C_6_ aniline-tagged CS disaccharide internal standard mixture (20pmole each) and 2 μl LC-MS grade water. CS disaccharides were analyzed by LTQ Orbitrap Discovery (Thermo Scientific) mass spectrometry attached with liquid chromatography system (Ultimate3000, Thermo-Dionex). All disaccharides were separated on a reverse phase C18 column (1mm x 150mm, 5µm, Targa, Higgins Analytical) using buffer containing dibutyl amine as ion-pairing reagent, followed by detection of disaccharides on mass spectrometer in negative ionization mode. The chromatographic steps were: 100% buffer-A (8mM acetic acid, 5mM DBA) for 10 min followed by stepwise increase of buffer-B amount; 17% buffer-B (70% methanol, 8 mM acetic acid, 5 mM DBA) for 10 min; 32% buffer B for 15 min; 40% buffer B for 15 min; 50% buffer B for 15min; 60% buffer B for 15 min; and 100% buffer A for 10 min. Flow rate was kept at 50µL per min.

### Calculation of ECM sulfate lost after chondroitinase digestion

In order to contextualize the Cl^-^_o_ increase we observe after chondroitinase treatment of OHSCs, we used our HPLC-derived data and approximated OHSC volume to derive a reasonable expected Cl^-^_o_ value for comparison. HPLC data was obtained from collected chondroitinase (or sham) digestion aCSF of a known volume. Chondroitin disaccharide concentration and sulfate content were found analytically, and a total chondroitin and total sulfate amount were calculated from mean values of chondroitin sulfate content (0.2mg/mL), sample volume (0.725mL), sulfation ratio (0.95 sulfate per disaccharide), and a monosulfated disaccharide molar mass of 481.4g/moL. Given mean OHSC thickness of 266μm and typical OHSC dimensions of 3×1.5mm, we calculate the best approximation of extracellular volume (20% of total slice volume) to be 0.293mL. The organic sulfate content lost from total ECS volume comes out to 1.65mM when assuming full-depth uniform chondroitinase digestion, or 8.8mM assuming depth-dependent digestion affecting only 50μm nearest the chondroitinase reservoir (i.e., nearest the perfusate). For comparison to measured Cl^-^_o_, we assume the molar relationship between sulfate lost and chloride gained to be 1:1.

### Image processing in Matlab

ABP-dextran fluorescence lifetime is converted to Cl^-^_o_ as described above. Composite Cl^-^_o_ images, depicting Cl^-^_o_ values via colormap while incorporating structural data through brightness adjustment, were produced using custom Matlab routines that apply normalized intensity data to Cl^-^_o_ images in a manner similar to commercial FLIM software. Note that intensity images were generated by summation of photons for each pixel across 32 time bins after excitation, resulting in a higher intensity background than is typical in standard two-photon images.

#### Pixel exclusion

We sought to perform as little processing of either intensity or lifetime data as possible, using statistical methods to objectively exclude pixels greater than three standard deviations from the mean of either property. Any remaining pixels reflecting a fluorescence lifetime outside the calibrated range (0-150mM Cl^-^), as happens in pixels with poor data quality typically due to low signal, were also excluded. More specifically, pixels were excluded if ABP- dextran lifetimes were in excess of the mean at 0mM chloride, or if lifetimes were shorter than twice the standard deviation less than the mean at 150mM (calculated from calibration data). This yielded total objective exclusion of 0.1-0.5% of pixels in a given image, which were set to black. Subjective exclusions were done either by manually drawn regions of interest (ROI) to exclude areas of perfusate at the margins of OHSCs or large blood vessels *in vivo*, or else by image-specific thresholding to exclude low-intensity areas coinciding with apparent somatic silhouettes. All exclusions were performed prior to analysis of extracellular Cl^-^_o_. To ensure that somatic silhouette-defined pixels, in particular, were not a subpopulation of pixels possibly reflecting ABP-dextran in an atypical compartment (e.g., the intracellular space), multiple analyses of excluded versus included pixels were performed to confirm faithful extracellular ABP-dextran signal and appropriateness of pixel exclusion (see Results).

#### Neuronal tracing analysis

Where somatic borders were traced for analysis, these were not dependent upon thresholding but were traced manually where neurons within the focal plane could be confidently identified based upon morphology. Traces followed the exterior aspect of the visually apparent border between bright and dim signal (see Results). Data was linearized and is presented as the pixelwise change in Cl^-^_o_ versus linearized distance, the latter calculated from pixel centerpoints. Comparison control data is derived by applying the same traces over calibration solution images, in which stochastic pixel to pixel differences are isolated (e.g., machine noise, Brownian motion artifact, etc). *In vivo* analysis was optimized for more superficial L1 cortical imaging and demonstration of ABP-dextran exclusion from vasculature and intracellular space, but a limited number of L2/3 neurons were also identified *post hoc* by their silhouettes formed through ABP-dextran exclusion within deeper fields. Morphological identification at this suboptimal optical zoom was only possible where pyramidal neurons were well within the chosen focal plane, producing demonstrably pyramidal silhouettes. This yielded a limited dataset on which we repeated perineuronal tracing analysis as an initial *in vivo* conformation of our better-optimized *in vitro* studies.

#### Regions of interest (ROIs) selection

ROIs are defined using intensity images for OHSCs either as a differentiation between tissue and surrounding bath, or this plus a regular division of the field (i.e., quarters or eighths) as indicated. ROI fields containing a majority of excluded pixels (e.g., perfusate) are excluded from analysis. ROIs are defined *in vivo* as cortical regions between anatomical landmarks, such as large vessels, which are easily identified through their exclusion of ABP-dextran. ROIs are then converted to masks and applied to fluorescence lifetime or Cl^-^_o_ images. ROIs in fields in which somatic silhouettes are evident are distinguished from ROIs that are too superficial to contain discernable silhouettes such as cellular (Layer 2/3) or acellular (Layer 1) cortical fields, respectively.

### Statistics

Statistical analysis was done with Matlab 2022a, with access to statistical toolbox functions. Individual tests used and *p*-values are detailed in figure legends. Corresponding *p*- value levels are indicated within figures by asterisks and listed in figure legends. Where sample sizes are small, we first determined whether we could assume normal distributions using the Jarque-Bera test (Matlab built-in function jbtest); where non-parametric tests were required, we utilized Mann-Whitney ranked sum or Kolmogorov-Smirnov tests as noted. *In vivo* statistical analysis of cortical Cl^-^_o_ was accomplished with ROI mean values, but because ROI size and number varies between groups we report mean+/-standard deviation of all analyzed pixels in addition to the ROI mean+/-standard error associated with ANOVA analysis. In the interest of transparency we elected to display violin plots of all pixels analyzed overlaid with ROI mean+/- standard error, using ANOVA analysis of the latter to determine significance of findings.

### Data availability statement

All data files used to generate the figures in the paper can be found on GitHub under KieranNormoyle/NeuronalChloride2023.

## Results

### Non-invasive measurement of extracellular chloride using ABP-dextran

ABP is a novel Cl- sensitive fluorophore compatible with <u>F</u>luorescence <u>L</u>ifetime <u>IM</u>aging (FLIM) that we have optimized to be bright, red-shifted, and sensitive to chloride concentrations through the range that would be encountered in in the extracellular space. We conjugated ABP to a 10,000 Dalton (10KDa) dextran resulting in an amine-bonded moiety (Figure 1A) that is resistant to pH changes in the physiological range (Normoyle *et al*., 2024) and is restricted to the extracellular space (see below). We validated the ability of ABP-dextran to reliably report chloride concentration by correlating the shortening of ABP-dextran fluorescence lifetime to chloride concentration (Figure 1B) through the Stern-Volmer relationship (Figure 1C), generating a *k_SV_* of 15.9M^-1^ for ABP-dextran used in these experiments. The fluorescence lifetime of ABP-dextran is not dependent upon the concentration of ABP-dextran, only the concentration of chloride.

**Figure 1.**
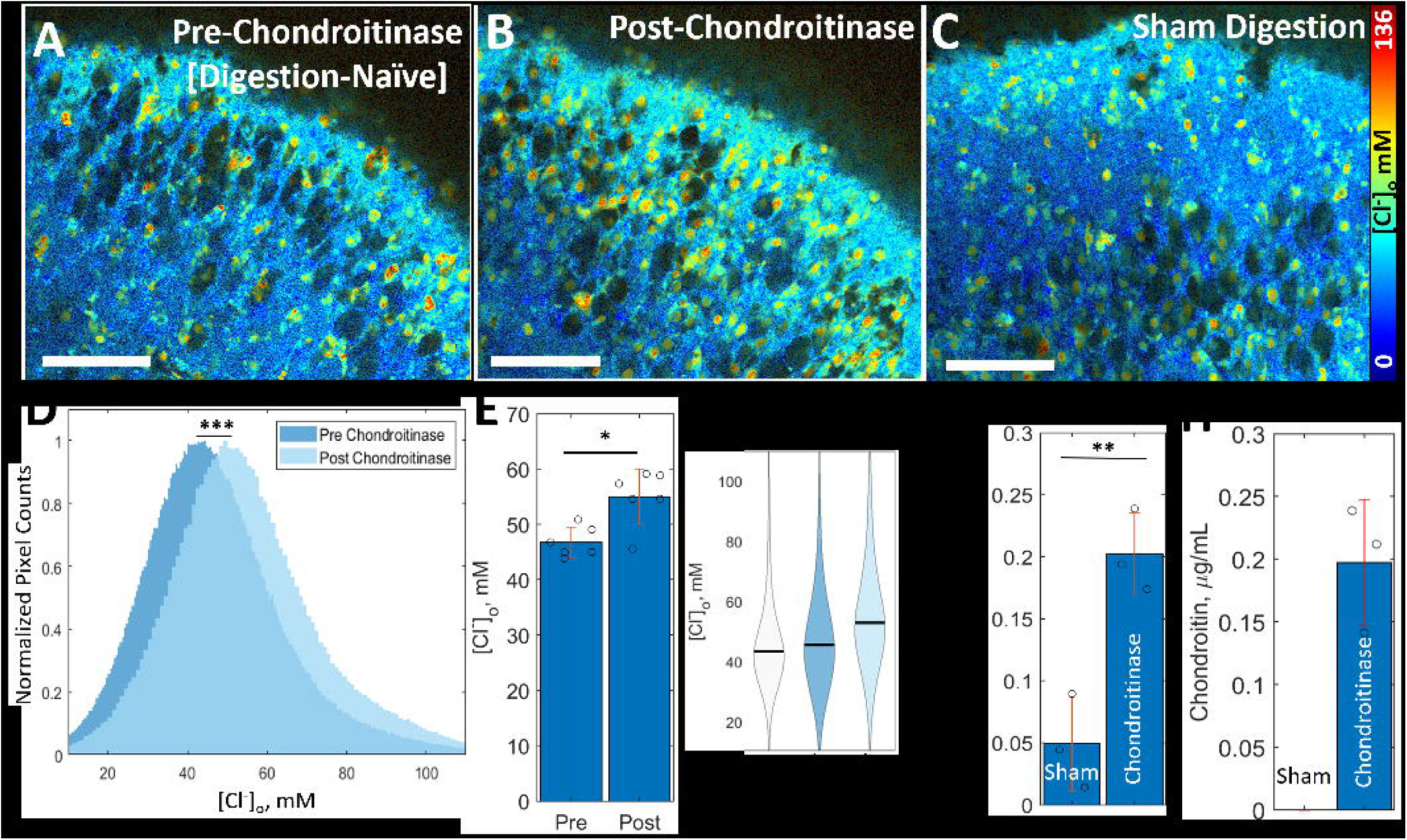
Characterization and Calibration of Chloride-Sensitive Fluorophore ABP. ***A)*** N- (**<u>a</u>**mmonium)-**<u>b</u>**utyl **<u>p</u>**henanthridine (ABP) is synthesized in-house and conjugated to activated 10KDa dextran, a glucose polymer, at a molar ratio of 4.4. ***B)*** Representative calibration of ABP- dextran fluorescence lifetime versus Cl^-^ demonstrating ABP-dextran sensitivity to chloride-induced quenching. Calibration of ABP-dextran fluorescence lifetime vs Cl^-^_o_ is displayed as a violin plot with mean +/- standard deviation overlaid. ABP-dextran lifetime at 0mM chloride (*τ_o_*) is extrapolated using a best-fit approach (see Methods). ***C)*** Stern-Volmer plot demonstrates a linear relationship between ABP-dextran fluorescence lifetime and concentration of chloride and was used to convert lifetimes to chloride concentrations. The ratio of unquenched ABP-dextran fluorescence lifetime (*τ_o_*) to ABP-dextran lifetime at a given concentration of quencher (*τ*) is plotted versus concentration of quencher (*τ_o_/τ* vs [Q^-^]). The slope of the linear regression is the Stern-Volmer constant (*k_SV_*) which allows conversion of ABP-dextran lifetime to concentration of quencher, specifically Cl^-^ (blue circles / black line; *k_SV_* = 15.9M^-1^). Four biologically relevant anions were found to have no experimentally relevant effect on ABP-dextran fluorescence lifetime: bicarbonate (*k_SV_* = 1.21M^-1^), gluconate (*k_SV_* = 0.70M^-1^), methyl sulfonate (*k_SV_* = 0.18M^-1^), and phosphate (*k_SV_* = 0.11M^-1^). Gluconate was subsequently used as a chloride substitute in non-standard aCSF formulations used for calibrations.

Chloride concentration also affects ABP-dextran intensity, such that high chloride concentrations not only shorten the emission lifetime but also reduce the intensity of emission. However, the concentration of chloride can be calculated (independently of the local ABP- dextran concentration) as long as sufficient photons are collected to calculate fluorescence lifetime in a given pixel.

### Sulfated agarose gels as an *in vitro* model of perineuronal space

We recently observed that differing concentrations of chloride juxtaposed to one another were stable on time scales of at least 100 minutes due to the presence of immobile anionic charges present in the neuronal cytoplasm (Rahmati *et al*., 2021), an effect known as Donnan exclusion (Marinsky, 1985; Helfferich, 1995; Fatin-Rouge *et al*., 2003; Epsztein *et al*., 2018; Gao *et al*., 2022). We tested whether Donnan exclusion was also operative in the extracellular space as a consequence of the abundant fixed negative charges present on extracellular glycolipids and glycoproteins (Schnaar *et al*., 2014), focusing in particular on the sulfate moieties of the glycosaminoglycans that comprise the bulk of the extracellular matrix in the brain (Karamanos *et al*., 2021). This matrix is most frequently visualized as the perineuronal lattice-like structures around all neurons and first described by Golgi in 1898 (Golgi, 1898; Celio *et al*., 1998). More recently the term ‘perineuronal nets’ has been applied to a specific extracellular structural motif associated with some interneurons and recognized by specific agents such as *wisteria floribunda* agglutinin (WFA); however, perineuronal space contains a ubiquitous, if variable (Deepa *et al*., 2006), abundance of negatively charged moieties (Schnaar *et al*., 2014) and Donnan exclusionary effects remain ubiquitous even as specific perineuronal environments vary among neuron subtypes. The process of Donnan exclusion does not require an ongoing energy input, although this issue has been contested (Glykys *et al*., 2014b; Voipio *et al*., 2014). We therefore modeled the brain’s extracellular matrix using a reduced system to remove the possibility of active buffering of Cl^-^_o_ by astrocytes (Egawa *et al*., 2013) or other cellular or vascular elements. We used gels of sulfated agarose to mimic the sulfated glycosaminoglycans such as the chondroitin sulfate-decorated aggrecan family members that are prominent in the perineuronal matrix. Agarose is a nominally sulfate-free linear glycopolymer that is a major component of agar (Figure 2B; bars 1 and 2) that was used as a chemically homogenous starting material for covalent sulfate linkage. We optimized the method of Fuse and Suzuki (Fuse & Toshiya, 1975) to balance the gelatinizing properties of agarose with the dispersive, hydrophilic Donnan effects arising from incorporation of the charged sulfate groups (Gregor, 1951; Helfferich, 1995). We obtained a sulfated agarose that could be used to cast gels with 50 mM effective sulfate concentration (Figure 2B, bar 3; 49.8+/-0.9mM sulfate across three batches [mean+/-SD]) covalently linked to the polymerized agarose lattice. Gels cast from sulfated agarose serve as a simple *in vitro* model of the perineuronal extracellular space, allowing us to investigate the ability of covalently linked anions attached to a three-dimensional matrix to displace mobile chloride ions.

**Figure 2.**
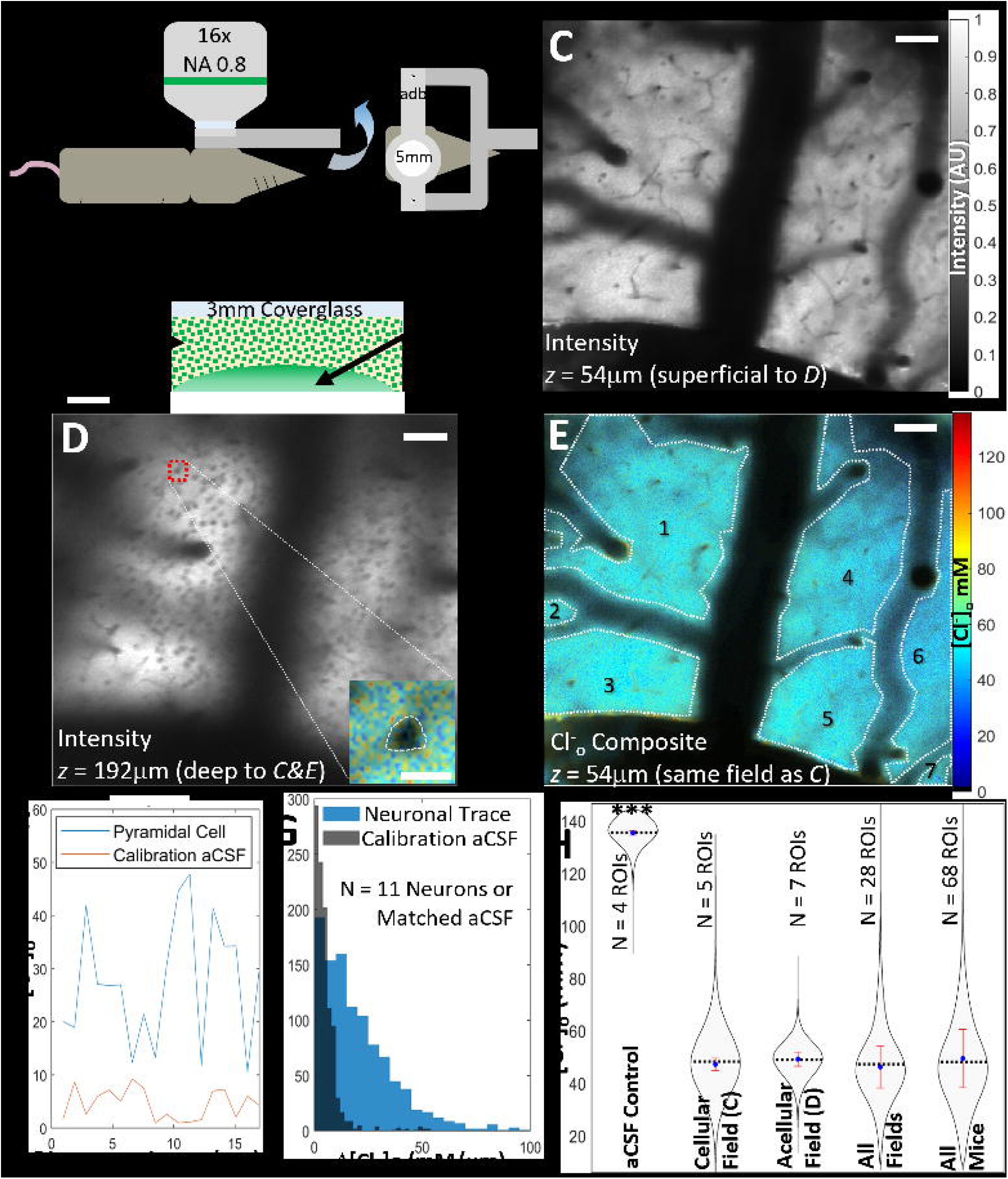
Effects of immobile sulfate on Cl^-^ in a volume of agarose gel. ***A)*** Experimental schematic demonstrating how polysaccharide gels, cast in aCSF separately, were equilibrated in aCSF after transfer to a common vessel. Gels were either equilibrated with aCSF containing ABP-dextran, processed for chemical analysis, or subjected to multiple investigations in tandem (see Methods). ***B)*** We covalently bound sulfate groups to algal biopolymer agarose and cast this as a gel in aCSF to model perineuronal interstitial matrix that is rich in chondroitin sulfate-decorated glycosaminoglycans. We compared sulfated agarose gel to native agar and agarose gels, achieving a 250-fold increase in sulfate content compared to commercially available agar (bar 3 vs bar 1, respectively; *p* < 0.001, Student’s paired *t*-test of at least three gel pieces’ mean of technical replicates; note discontinuous y-axis). Commercially available agarose, which is intended as a sulfate-free product, had no measurable sulfate content (bar 2) and subsequently was used as a control gel. ***C)*** Chloride concentration was reduced in sulfated agarose gels. Cl^-^ was measured using both a standard colorimetric assay (dark gray bars) and ABP-dextran FLIM (light gray bars) in aCSF control (leftmost pair), within non-sulfated agarose gel (center pair) and within sulfated agarose gel containing ≈50mM (49.2+/-0.289mM) covalently bound sulfate (rightmost pair). Measured chemically or using ABP-dextran FLIM, Cl^-^ in sulfated agarose gels differed significantly from aCSF (*p* = 0.00264 and *p* < 0.001, respectively) and non-sulfated agarose gels (*p* = 0.00354 and *p* < 0.001, respectively), while Cl^-^ measured in non-sulfated agarose gels was not significantly different from aCSF when measured by chemical assay (*p* = 0.190) but registered a small but significant difference (3.5% increased Cl- relative to aCSF, *p* = 0.00328) when measured with ABP-dextran FLIM. Chemical assay and aCSF FLIM data points are averages of three technical replicants, while gel values are averaged measurements from each quadrant of a given gel, each compared using Student’s paired *t*-test. ***D)*** Significant chloride concentration differences between agarose gels with 50mM covalently bound sulfate (dark blue) and non-sulfated agarose gels (light blue) do not equilibrate with time but instead are stable for as long as experimentally observed (> 3 weeks). ANOVA testing with Bonferroni correction demonstrates that Cl^-^ significantly differs between native agarose and sulfated agarose at all time points (overall and each pair *p* < 0.001) while neither agarose (each *p* = 1.0000) nor sulfated agarose gels (from left to right, *p* = 1.0000; *p* = 0.2119; *p* = 0.1129) change significantly with time. Cl- measurements from each quadrant of a given gel are averaged to generate each data point. ***E)*** Demonstration of stable juxtaposition of significantly different chloride concentrations at equilibrium. Separately cast sulfated and non-sulfated gel pieces are placed in a single vessel as described in ***A*** and equilibrated in common aCSF with ABP-dextran. Sulfated agarose gels (dark blue) again contain significantly lower Cl^-^ than do non-sulfated agarose gels (light blue; *p* < 0.001) or perfusate aCSF (gray; *p* < 0.001) in this representative experiment, while Cl^-^ within non-sulfated agarose gels is again not significantly different than perfusate aCSF (*p* = 0.0647)., Fields from each quadrant of each gel co-incubated in a single vessel with shared aCSF are analyzed using Student’s paired *t*-test. *Statistical significance indicated on graph for p-values <0.05 (*); <0.01 (**); <0.001 (***); “ns” indicates “not significant”. N = 3 except for **D**, where N = 4. Error bars reflect standard deviation from the mean. Use of parametric tests confirmed appropriate via Jarque-Bera test (see Methods)*.

We measured the chloride concentrations within non-sulfated agarose gels vs. sulfated agarose gels, each equilibrated in aCSF, and compared these to the chloride concentration in the surrounding aCSF using either a standard colorimetric iron(III) thiocyanate assay (Figure 2C, dark gray bars) or ABP-dextran FLIM imaging (Figure 2C, light gray bars). Agarose gels with fixed anions in the form of covalently bound sulfate had significantly decreased chloride concentrations versus both control aCSF and non-sulfated agarose gels (Figure 2C, right pair vs left and center pairs, respectively) measured either with the standard chemical assay (dark gray bars; *p* = 0.00264 and *p* = 0.00354 by Student’s paired *t*-test, respectively) or ABP-dextran FLIM (light gray bars; *p* < 0.001 by Student’s paired *t*-test for either comparison). We measured a small chloride increase within non-sulfated agarose gels vs control aCSF in either assay, representing an insignificant increase when using the standard chemical assay (8.05+/-6.14mM, mean+/-standard deviation; *p* = 0.190 by Student’s paired *t*-test) and a smaller yet statistically significant increase when measured using ABP-dextran FLIM (4.95+/-0.51mM, mean+/- standard deviation; *p* = 0.00328 by Student’s paired *t*-test). Later ABP-dextran FLIM experiments comparing aCSF perfusate to non-sulfated gels show no such difference (see below and Figure 2E). This Cl^-^ increase is curious but is quantitatively small and qualitatively distinct from the Cl^-^ decrease we see in sulfated agarose gels (3.5% increase from non-sulfated gel vs. aCSF control compared to a 40.7% decrease from sulfated agarose gel vs. aCSF control, respectively), meaning that any effect would be a small overestimation of Cl^-^ within sulfated agarose gels. The overall agreement of results derived from both assays lends confidence to the reliable performance of ABP-dextran FLIM versus standard chemical colorimetric methods of Cl^-^ measurement.

Next we placed separately cast non-sulfated and sulfated gels adjacent to each other and equilibrated in the same aCSF bath with ABP-dextran (Figure 2D; schematic in Figure 2A). ABP-dextran FLIM measurements of chloride concentration within the non-sulfated and sulfated gels demonstrated significant chloride concentration differences between gels that had no fixed anionic charge to displace mobile chloride (non-sulfated agarose; Figure 2D light blue bars) vs. gels that had covalently bound sulfate groups (sulfated agarose; Figure 2D dark blue bars). The chloride concentrations were inversely proportional to the sulfate concentration in the gel, were significantly lower for sulfated gels at each time point (each *p* < 0.001), and were stable for over three weeks (agarose gel comparisons: each *p* = 1.0000; sulfate gel comparisons: 2-4hours vs 24hours *p* = 1.0000, 3weeks vs 2-4 hours and 24 hours *p* = 0.211 and *p* = 0.113, respectively; ANOVA with Bonferroni correction). These findings support the displacement of mobile chloride ions by the sulfates fixed to the gel (Figure 2D) by Donnan exclusion. We next imaged both non-sulfated and sulfated gels immediately juxtaposed to one another, along with aCSF perfusate, in the same field (Figure 2E) and demonstrated that the presence of sulfate groups covalently bound within the polymerized gel results in a significantly decreased chloride concentration relative to both non-sulfated gels and aCSF perfusate (Figure 2E, bar 3 vs bar 2 and bar 1, respectively; each *p* < 0.001 by Student’s paired *t*-test). Non-sulfated agarose gels demonstrated no difference versus aCSF perfusate (*p* = 0.0647 by Student’s paired *t*-test).

### Demonstration of chloride displacement in living systems

To test whether Donnan exclusion is also operative in the extracellular matrix of the brain, we studied mouse hippocampal organotypic slice cultures. We tested whether extracellular perineuronal fixed anions, such as the chondroitin sulfate moieties of aggrecan family proteoglycans (Frantz *et al*., 2010; Karamanos *et al*., 2021), displace chloride in this living system. After incubation with ABP- dextran and equilibration with aCSF containing ABP-dextran and 136mM chloride, we measured extracellular chloride concentration using ABP-dextran FLIM and found that Cl^-^_o_ was unexpectedly low and heterogenous (46.7 +/- 18.8 mM, calculated from 25.8×10^6^ pixels across 32 fields of 7 slices). Representative fields are shown in Figure 3A (ABP-dextran intensity) and Figure 3B (Cl^-^_o_ composite image, quantified in Figure 3C). Paired analysis of slices pre- and post-ABP-dextran incubation confirmed that autofluorescence signals in pixels in pre-ABP images did not contribute to the unexpectedly low values of Cl^-^_o_ obtained from post-ABP images. This is because pre-ABP pixels that had both sufficient photon count to allow lifetime calculation and lifetimes within the calibrated range of ABP-dextran had short lifetimes whose effect would be to increase the reported Cl^-^_o_, not decrease it.

**Figure 3.**
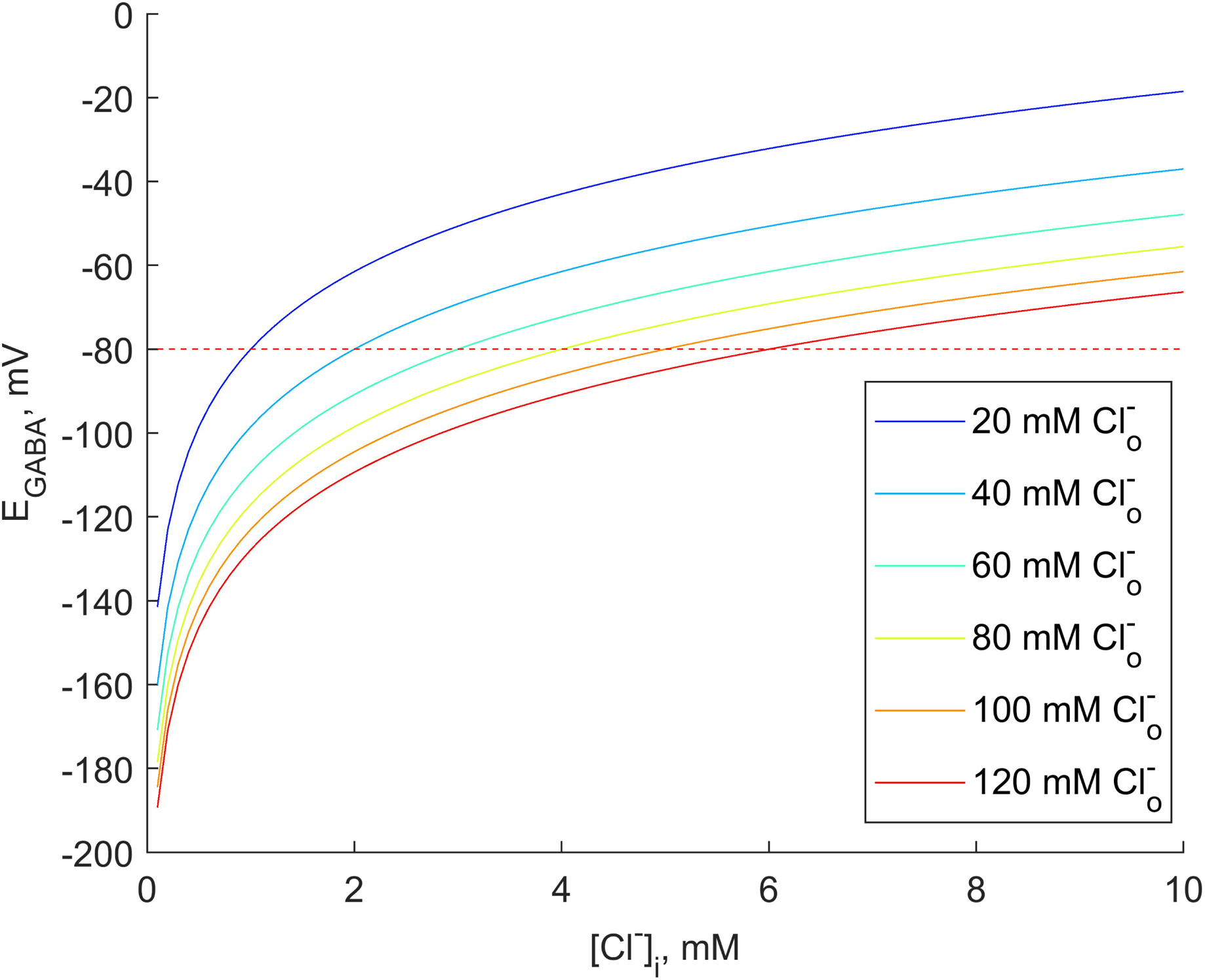
Cl^-^ in the extracellular space is not uniform and is considerably lower than aCSF perfusate. ***A)*** High-magnification OHSC intensity image of ABP-dextran excluded from cell bodies in the pyramidal cell layer (CA1; scale bar 20μm). ***B)*** Chloride-composite image again demonstrating the exclusion of ABP-dextran from intracellular spaces as well as the non-uniformity of Cl^-^_o_ in the pyramidal cell layer of a representative OHSC. Representative pyramidal cell visible just above and left of image center, identified by ABP-dextran exclusion and morphology of resulting silhouette, is one of 6-10 neurons per field deemed to have a regular and nearly-continuous border analyzed as described in ***D&E*** (scale bar 20μm). ***C)*** Histogram of full field Cl^-^_o_ in OHSC pictured in ***B***, demonstrating non-uniformity visually apparent in ***B***. Notably, Cl^-^_o_ = 44.5+/-15.9mM despite perfusate [Cl^-^] = 136mM (across all depths of this slice Cl^-^_o_ = 49.7+/-18.5mM; mean +/- SD). ***D)*** Change in chloride (Δ[Cl^-^]_o_) plotted against distance in the perineuronal space (blue line) or the same trace overlaid onto a calibration aCSF FLIM image (red line) corresponding to the perineuronal trace in ***B***. Δ[Cl^-^]_o_ of calibration aCSF (red) encapsulates machine and experimental noise, and Δ[Cl^-^]_o_ in the perineuronal space (blue) exceeds that of control in this representative neuron. ***E)*** Perineuronal Δ[Cl^-^]_o_ per micron calculated from many neurons’ circumferential traces demonstrates that the variability of perineuronal Cl^-^_o_ in living *in vitro* system exceeds the variability of control aCSF (72 neurons versus matched calibration aCSF controls from depths of 20-100μm in this representative OHSC; p < 0.001, by Kolmogorov-Smirnov test; *N* = >26,000 pixel-wise measurements across 72 perineuronal traces). *Cl^-^_o_ image rendering and presentation details can be found in Methods and in subsequent figures. Scale bars A&B: 20μm*.

Perineuronal traces along the outer edge of neuronal silhouettes (Figure 3B, white dashes) were used to analyze the spatial change in Cl^-^_o_ in the extracellular space immediately adjacent to the neuronal cytoplasmic membrane. Pyramidal neuron silhouettes were identified morphologically and pixels underlying the extracellular margin (i.e., perineuronal) traces were linearized for analysis. We compared the size of the effects of fixed anions on local Cl^-^_o_ to Brownian and machine noise arising from FLIM of ABP-dextran in bulk solutions with similar mean Cl^-^_o_. The pixel-to-pixel change in Cl^-^_o_ (ΔCl^-^_o_) versus distance in the presence of fixed anions in the perineuronal space (Figure 3D, blue) was much larger than the ΔCl^-^_o_ versus distance in the absence of fixed anions in bulk solution containing ABP-dextran and a single concentration of Cl^-^ (Figure 3D, red). Figure 3E shows similar data for a larger cohort of perineuronal tracings and their corresponding aCSF solution counterparts (72 neurons traced from 6 OHSCs). We found a consistently increased probability of greater ΔCl^-^_o_ per micron in the extracellular space where fixed extracellular anions were present compared to the perfusate aCSF (Figure 3E, blue vs red).

We further analyzed dim pixels typical of the neuronal silhouettes, which had not already been statistically excluded and displayed as black. These dim pixels retain their Cl^-^_o_ colormap values but are often so dim they are difficult to appreciate visually (e.g., Figure 3B). They are included in presented images in the interest of transparency but were excluded from Cl^-^_o_ analysis after applying an empirically derived threshold value that best approximates the clearest silhouettes in each image. We reasoned that these pixels represented photons originating from extracellular ABP-dextran above or below the focal plane, likely present due to the higher-background FLIM-derived intensity images (Figure 4A) which represent a summation of all photons in each pixel. Because Cl^-^_o_ is derived from the intrinsic, intensity-independent property of fluorescence lifetime, structural context is lost in Cl^-^_o_ images (Figure 4B) and must be reestablished by incorporation of intensity information into a composite image (Figure 4C). This is not unique to Cl imaging; commercial FLIM software creates intensity-lifetime composite images in a similar manner. In our images we first converted from fluorescence lifetime to Cl^-^_o_ to allow the use of a linear colormap for readers’ ease of interpretation (see Methods).

**Figure 4.**
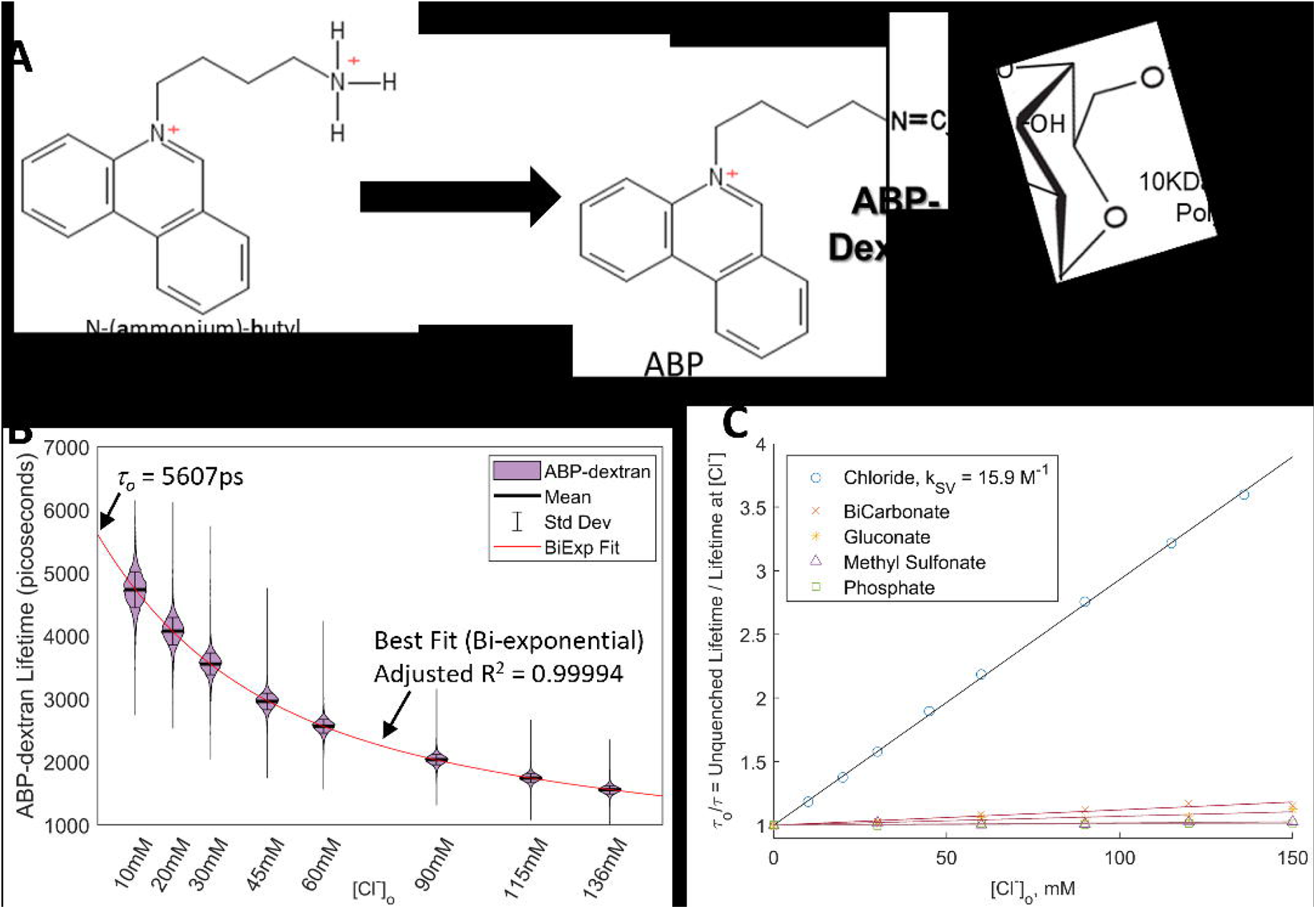
Generation of composite chloride images and cell body masking for pixel quantification. ***A)*** ABP-dextran intensity data of an OHSC, derived by taking the total number of photons detected in each pixel throughout the acquisition period, is used to inform on tissue structure. Image series ***A-C*** corresponds to Figure 3B and its traced neuron example. Neuron silhouette traced in ***B&C*** is indicated with a white asterisk. ***B)*** Cl^-^_o_ calculated from ABP-dextran lifetime. The loss of discernible structure is due to the fact that the intensity of the ABP-dextran signal has no effect on the Cl^-^_o_ calculated from fluorescence lifetime (see Methods). Areas corresponding to cell body silhouettes in ***A*** have notable pixelation with relatively high apparent Cl^-^_o_ values interspersed with black pixels, the latter indicating statistical exclusion due to low signal intensity (see Methods). ***C)*** In order to observe Cl^-^_o_ with spatial context restored, we use intensity data to inform the brightness of the raw chloride data in ***B***, generating a composite image in which the color corresponds to Cl^-^_o_ and brightness is attenuated based on the intensity of the signal in each pixel (see Methods). The most popular proprietary FLIM software packages use similar methods of composite data display. Note that prior to Cl^-^_o_ analysis a further intensity-based thresholding step excludes pixels from inside somatic silhouettes in the focal plane, and pixels not corresponding to the slice (i.e., perfusate, top left) are also excluded by manually drawn ROI (masks not shown). ***D)*** Histogram of Cl^-^_o_ in all analyzed pixels (blue) versus pixels excluded through thresholding step, principally marking somatic silhouettes (gray) in this representative OHSC, demonstrating no secondary peak at typical Cl^-^_i_ values (i.e., =< 15mM). Further, this range of low Cl^-^_o_ values is more likely excluded (gray) than included (blue) in our analysis (indicated by gray arrow). ***E&F)*** To further verify the cellular exclusion of ABP-dextran, we looked specifically at the margins of well-defined somatic silhouettes, such as the example marked across ***A-C***, to search for signs of intracellular ABP-dextran leakage. We compared perineuronal pixels up to 1mm outside neurons to those pixels which lie inside the somatic margins of 72 neurons but were not excluded prior to thresholding (statistically excluded pixels are not considered; see Methods). Consistent with out-of-focus extracellular ABP-dextran, we found similar calculated Cl^-^_o_ values between the two groups (***E***) with the expected wider variance in the less intense intracellular signal (***F***; see text). *Scale bar: **A-C** 50μm*.

The dim, excluded pixels occurring in neuronal silhouettes offered an opportunity to assess whether they were in fact caused by out-of-focus extracellular ABP-dextran, consistent with recent super-resolution imaging of the extracellular space utilizing a 10KDa dextran-linked fluorophore in addition to unconjugated fluorophore (Tønnesen *et al*., 2018). If true, this would produce a similar, if wider (Normoyle *et al*., 2024), distribution of Cl^-^_o_ values. Conversely, if the dim, excluded pixels were autofluorescent in origin or were reflective of low Cl^-^_i_ reported by ABP-dextran leaked into neurons, then we would find a right-shifted, high-Cl^-^_o_ distribution or a left-shifted, low-Cl^-^_o_ distribution, respectively.

### Verification of ABP-dextran extracellular compartmentalization

We used two approaches to assess the Cl values reported in the dimmest pixels. We first compared all pixels below the exclusion threshold to all pixels above the threshold. Only pixels outside the actual slice (i.e., perfusate pixels) and those pixels with lifetimes outside the calibrated range of ABP-dextran were not considered. We found that included and excluded pixels had histograms with similar characteristics, such as mean and mode, although excluded pixels were fewer in number and had a wider distribution (Figure 4D, blue vs gray). There is neither a high-Cl^-^_o_ peak indicative of autofluorescence nor is there a low-Cl^-^_o_ peak indicative of intracellular signal in either included pixels or excluded pixels. Notably, there are virtually no included pixels in the typically intracellular range (i.e., ≤15mM Cl^-^), and such pixels are far more likely to be excluded from analysis (Figure 4D, gray arrow). We conclude that intensity-excluded pixels are not a low- or high-Cl^-^_o_ subset, which would seem to preclude errant Cl^-^_i_ signal. However, we wanted to rule out any possible non-extracellular signal more specifically as most pixels at these resolutions are expected to contain a portion of both neuropil and extracellular space.

As a second means to evaluate the Cl values reported in the dimmest pixels, we assessed pixels within individual somatic silhouettes. Visually apparent somatic edges are ideal to compare extracellular ABP-dextran signal in the bright perineuronal exterior of silhouettes to ostensibly intracellular pixels inside the silhouette margins. The fact that the brightest of these pixels are located nearest the outer silhouette margin and the least intense pixels are located toward silhouette centers supports our hypothesis that this signal is extracellular in origin from outside the focal plane. Alternatively, if ABP-dextran leaked inside cells, despite its conjugation to 10KDa dextran, we would expect that such an intracellular signal would be brightest in the silhouettes formed by cells firmly within the focal plane (i.e., with the most ABP-containing cytoplasm in the focal plane). After the objective pixel exclusion step (intensity greater than three standard deviations from the mean, or lifetime outside the calibrated range), we compared Cl^-^_o_ in the perineuronal margin within one micron external to the traced border to those pixels inside the traced border. In this way our analysis was restricted to only those cells morphologically identified as pyramidal neurons adequately centered in the focal plane. Pixels from inside the silhouette are greater in number than perineuronal extracellular pixels in this analysis, yet we saw no subpopulation of intracellular chloride values (i.e., in the range of 15mM Cl^-^) that would be expected if this signal were originating from ‘leaked’ intracellular ABP-dextran (Figure 4E). Signal from the one micron external to the perineuronal perimeter is five times as bright on average as pixels from within the silhouette (Figure 4F), which helps explain why the excluded intracellular signal has a wider distribution of Cl^-^_o_ values (Normoyle *et al*., 2024). If no intracellular leakage of ABP-dextran can be detected even within in-focus neuronal somatic silhouettes, where such signal is expected to be strongest if it were present, one can infer that neuropil will not contribute any such signal in pixels containing both neuropil and extracellular space. While we look forward to future experiments with greater spatial resolution, we conclude that neither objectively excluded pixels nor those excluded by intensity thresholding are indicative of low- or high-Cl^-^_o_ subsets. Further, we provide ample evidence supporting the fidelity of ABP-dextran in its reporting of chloride exclusively in the extracellular space.

### Demonstration of chloride displacement by chondroitin sulfate

We then confirmed that fixed anions, including chondroitin sulfate glycoproteins, are a significant contributor to the observed difference between expected and measured Cl^-^_o_ by measuring Cl^-^_o_ before and after treatment with chondroitinase. Analogous to matrix metalloproteases, chondroitinase is a bacterial enzyme that cleaves the sulfated glycopolymer chondroitin. Chondroitinase treatment significantly increased Cl^-^_o_ in organotypic hippocampal slice cultures (Figure 5A&B; pre- and post-digestion, respectively). Cl^-^_o_ increased on average 8.3+/-3.3mM (mean+/-standard deviation) in freely diffusible extracellular chloride within the slice culture (Figure 5A vs 5B; quantified in 5D-F) and was significantly greater in chondroitinase digested slices than in sham digested slices (Figure 5B vs 5C; quantified in 5F). This increase in Cl^-^_o_ coincides with a significant decrease in the extracellular fixed anions within the slice culture. While chondroitin sulfate is targeted in this experiment, proteoglycans collectively make up less than 10% of the extracellular fixed charge; gangliosides and sulfatides make up the balance (Schnaar *et al*., 2014). The actual decrease in fixed anion concentration in the extracellular space was estimated from the concentration of chondroitin sulfate released into the perfusate after chondroitinase treatment (Figure 5G) and confirmed by HPLC analysis (Figure 5H). Quantitative estimates of fixed anionic charge in the perineuronal space are few in the absence of a quantitative fluorescent or histological sensor. Those present in the literature are either based on novel techniques (Marinsky, 1985) or else are extrapolated from non-neuronal tissues also rich in chondroitin sulfate (Urban *et al*., 1979; Chahine *et al*., 2005). We estimated the expected Cl^-^_o_ increase after chondroitinase digestion using HPLC-derived measurements of released chondroitin disaccharides and OHSC volume (see Methods). Careful *in vitro* studies of chondroitinase have concluded that only approximately 50% of total chondroitin sulfate is accessible to chondroitinase (Lin *et al*., 2008). This is likely not due to a diffusion restriction (Hrabetová *et al*., 2009) and more likely is related to enzyme activity, perhaps due to variance in local matrix structures. Our own empirical observations in OHSCs, in agreement with others’ published observations (e.g. Orlando and colleagues, especially Figure 5C therein) (Orlando *et al*., 2012) indicate that chondroitinase activity decreases with distance from its diffusive reservoir and full activity is restricted to an estimated 50μm; as such we only measure Cl^-^_o_ to a depth of 50μm. Rather than seek definitive conclusions about the enzymatic efficiency of chondroitinase at this time, we performed calculations for each case. If all digested chondroitin sulfate indeed came from only the most superficial 50μm of our OHSCs, the Cl^-^_o_ increase expected in this same superficial 50μm is calculated to be 8.8mM (see Methods). This agrees well with experimental mean Cl^-^_o_ values calculated from regular subfields of pre- and post-chondroitinase digestion, 8.8mM versus 8.3+/-3.3mM (Figure 5E), or when using the full field to calculate the minimum and maximum range of Cl^-^_o_ increases from 0.6-16.1mM (Figure 5D; pre-digestion mean minus standard deviation to post-digestion mean plus standard deviation). Further, we find that even if we assume chondroitin sulfate is lost uniformly from the full thickness of the OHSC and recalculate a smaller expected Cl^-^_o_ increase of 1.65mM, this still lies within the empirically measured range of 0.6-16.1mM. These observations are consistent with the expected inverse relationship between fixed extracellular anions such as sulfate and mobile Cl arising from Donnan exclusion. They are also consistent with the fact that chondroitin sulfate, while experimentally accessible, only accounts for a minority of total expected perineuronal fixed anionic charge despite being the dominant sGAG in the perineuronal space. Estimations employing bulk biochemical measurements from rat brain (Schnaar *et al*., 2014) suggest that the actual fixed anionic charge contribution from chondroitin sulfate is approximately 2mM equivalent charge (see Methods) in agreement with our findings and consistent with the electrophysiological findings of others (Klein *et al*., 2018). It would be advantageous to similarly estimate a predicted Cl^-^_o_ based upon local sulfate concentrations and compare this to empirical Cl^-^_o_ measurements similar to our sulfated agarose experiments (Figure 2E), but without direct and independent measurement of the extracellular sulfate content of OHSCs this is not currently feasible. In summary, the immobile ions associated with the extracellular matrix displace mobile anions such as chloride consistent with the well-established effects of Donnan equilibrium forces, such that Cl^-^_o_ is much lower and less uniform than would otherwise be expected.

**Figure 5.**
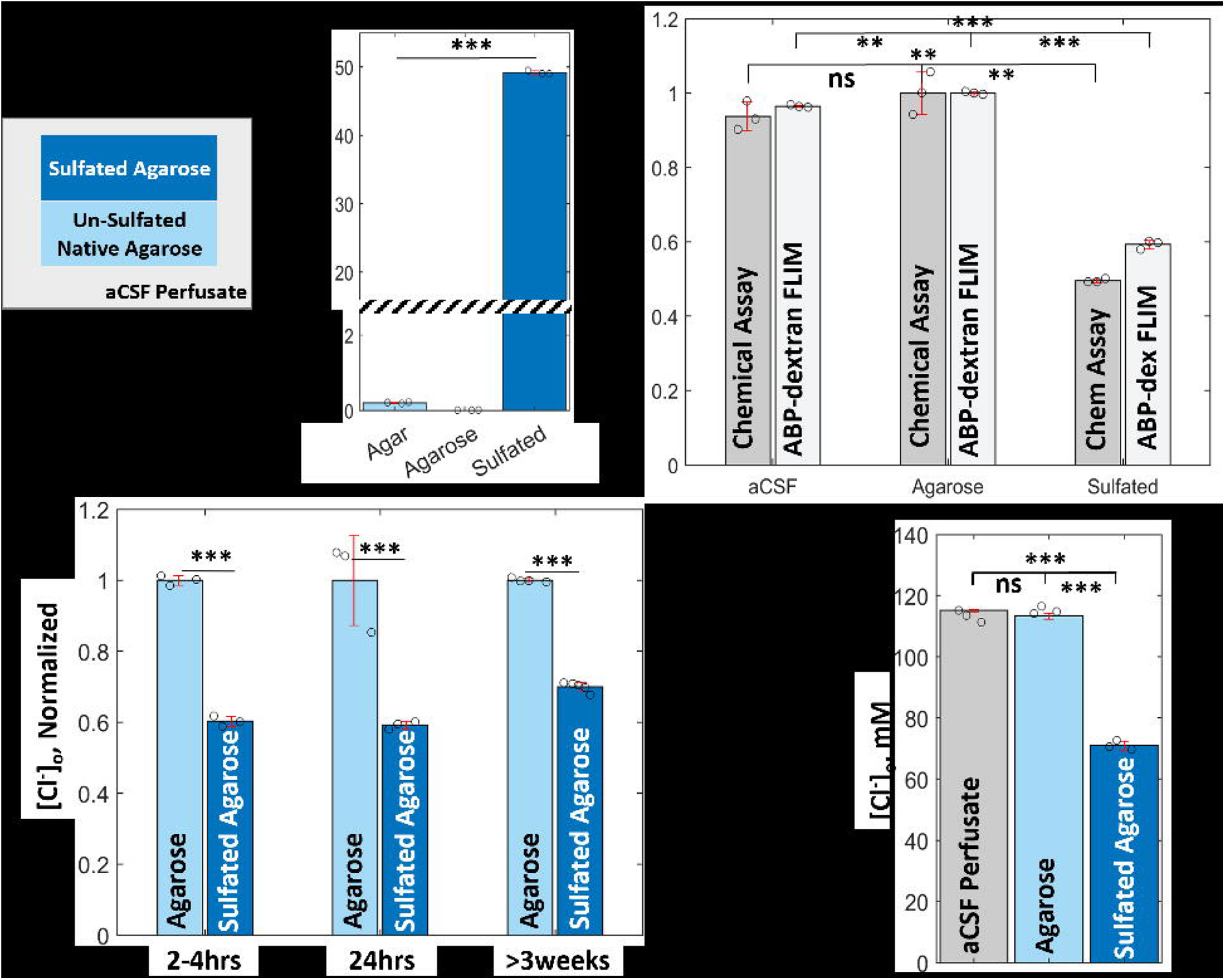
Chondroitinase treatment removes fixed perineuronal anions and attenuates Cl^-^_o_ displacement. Cl^-^_o_ composite images of representative mouse hippocampus slice cultures imaged in culture media (Cl^-^ ≈ 64mM) with 1mg/mL ABP-dextran (100μM ABP-Dextran) before (***A***) and after (***B***) treatment with bacterial MMP analog chondroitinase, or after sham digestion (***C***); shared colorbar for ***A-C*** at right. ***D)*** Histograms of [Cl^-^]_o_ corresponding to ***A&B*** demonstrate pre-chondroitinase digestion (dark blue) and post-chondroitinase digestion (light blue) [Cl^-^]_o_ populations of pixels are significantly different (*p* < 0.001 by Kolmogorov-Smirnov paired-sample test). ***E)*** When images from ***A&B*** are broken into regularly shaped regions of interest and analyzed as paired groups, pre-chondroitinase and post-chondroitinase are again significantly different (46.7+/-2.7mM vs 55.0+/- 5.0mM, respectively, mean+/-SD; *p* = 0.0152 by Wilcoxon rank sum test; *N* = 6). ***F)*** Violin plot of data from ***A&B*** (dark and light blue, respectively) vs grouped sham-digested slice data (gray) demonstrating that Cl^-^_o_ increases significantly after chondroitin sulfate is removed by digestion, whether compared to pre-digestion of the same slice or a sham digestion (each *p* < 0.001 by Kolmogorov-Smirnov paired-sample test). ***G)*** Measurement of glycosaminoglycans, including chondroitin sulfate, released into the perfusate after OHSCs are treated with chondroitinase (vs sham digestion) using a commercial colorimetric assay. A significantly higher concentration of polyanionic glycosaminoglycans are found in the extracellular fluid after chondroitinase digestion (*p* = 0.00632 by Student’s paired *t*- test; *N* = 3), consistent with the release of chondroitin sulfate from extracellular matrix within slice cultures and a resultant increase in Cl^-^_o_. ***H)*** HPLC analysis confirms the presence of chondroitin sulfate released into digestion buffer after chondroitinase treatment (mean = 0.20μg/mL; *N* = 3); there was no detectable chondroitin sulfate in sham digests, negating formal significance testing. ***A-C*** *scale bars 50μm; all slice cultures were DIV14-21 at time of experiment; cultures were grown directly on glass with the exception of **G** where cultures were grown on membrane inserts (see Methods); significance values noted graphically for p-values <0.05 (*); <0.01 (**); <0.001 (***); “ns” indicates “not significant”. Use of parametric tests confirmed appropriate via Jarque-Bera test (see Methods). Histograms in D&F include at minimum 78,163 pixel-wise chloride measurements. Note that data contained in **D-F** labeled ‘pre’ and ‘post’ are derived from OHSCs in **A** and **B**, respectively*.

### Chloride displacement in the perineuronal space *in vivo*

To test whether Donnan exclusion of Cl^-^_o_ by sulfated GAGs and other fixed anions was significant *in vivo*, we performed live 2 photon imaging of P26-30 mice via a cortical window (Figure 6A). Direct application of high concentrations of ABP-dextran into the non-sulfated agarose overlying the neocortex (Figure 6B) resulted in a robust ABP fluorescence signal from the extracellular space of the brain parenchyma, similar in intensity to 50μM ABP-dextran used *in vitro*. Control measurements were obtained from the agarose above the cortex, which had been brought up in aCSF (136mM Cl^-^) and allowed to equilibrate with *in vivo* CSF. Notable exclusion of ABP-dextran fluorescence emission from blood vessels (Figure 6C,D&E) and cell bodies (Figure 6D) was observed. Areas of locally elevated Cl^-^_o_ and overall bright signal seen in OHSCs (Figures 3-5) were absent *in vivo* (Figure 6), suggesting these features are specific to *in vitro* preparations. At several microns in size, we reason the areas seen in OHSCs are most likely damaged areas where intracellular space makes up less than the typical 80% of volume allowing ABP-dextran access to a greater proportion of volume, so that the signal is bright despite quenching by the high Cl^-^_o_ in these spaces. ECM damage would also be likely in such circumstances and would be expected to result in a loss of extracellular immobile anions, limiting the Donnan exclusionary force and leading to relatively high concentrations of Cl^-^_o_. This combination would be expected to result in a paradoxically bright, short-lifetime ABP-dextran signal yielding an unusually intense high Cl^-^_o_ area. Given the absence of these features *in vivo*, we speculate that these areas are the result of slicing trauma but longitudinal studies will be needed to adequately categorize them. Regions of interest (ROIs) were defined using intensity images before rendering composite images so that only structural information was considered; an example intensity and composite image pair can be seen in Figure 6C&E. Perineuronal tracing analysis was repeated for the limited number of layer 2/3 neurons we could identify by morphology and confirmed that perineuronal Δ[Cl^-^]_o_ per micron *in vivo* is similar to *in vitro* findings (Figure 6D inset, F&G; compare to Figure 3D&E). Cortical Cl^-^_o_ was less than half the chloride concentration measured in the overlying agarose (48.1 +/- 17.0mM, calculated from 68 ROIs and 4 animals, versus 135.4 +/- 5.0mM in agarose between cortex and coverglass). Cl^-^_o_ in cellular and acellular fields displayed in Figure 6C,D&F were not significantly different from each other (Figure 6E entries 2&3, respectively), nor were either significantly different from all ROIs collected from this animal (entry 4) or from all animals (entry 5; each p = 1.0000 by ANOVA with Bonferroni correction), while all were significantly different when compared to aCSF measured above the cortex (each p < 0.001 by ANOVA with Bonferroni correction). Regional mean Cl^-^_o_ values were approximately 45-55mM (Figure 6E) but there was notable spatial variation between regions and even within regions, similar to *in vitro* OHSC results and consistent with local Donnan exclusion of Cl^-^_o_ by local variation in fixed anions such as sulfated GAGs. Future experiments will be optimized for higher resolution imaging, but our results suggest that the well-established heterogeneity of sulfation patterns around neurons represent a means by which neurons and supporting cells can indirectly modulate Cl^-^_o_ through Donnan forces in a manner similar to how cytoskeletal components, in particular actin, have been recently shown to influence local intracellular Cl^-^_i_ (Rahmati *et al*., 2021).

**Figure 6.**
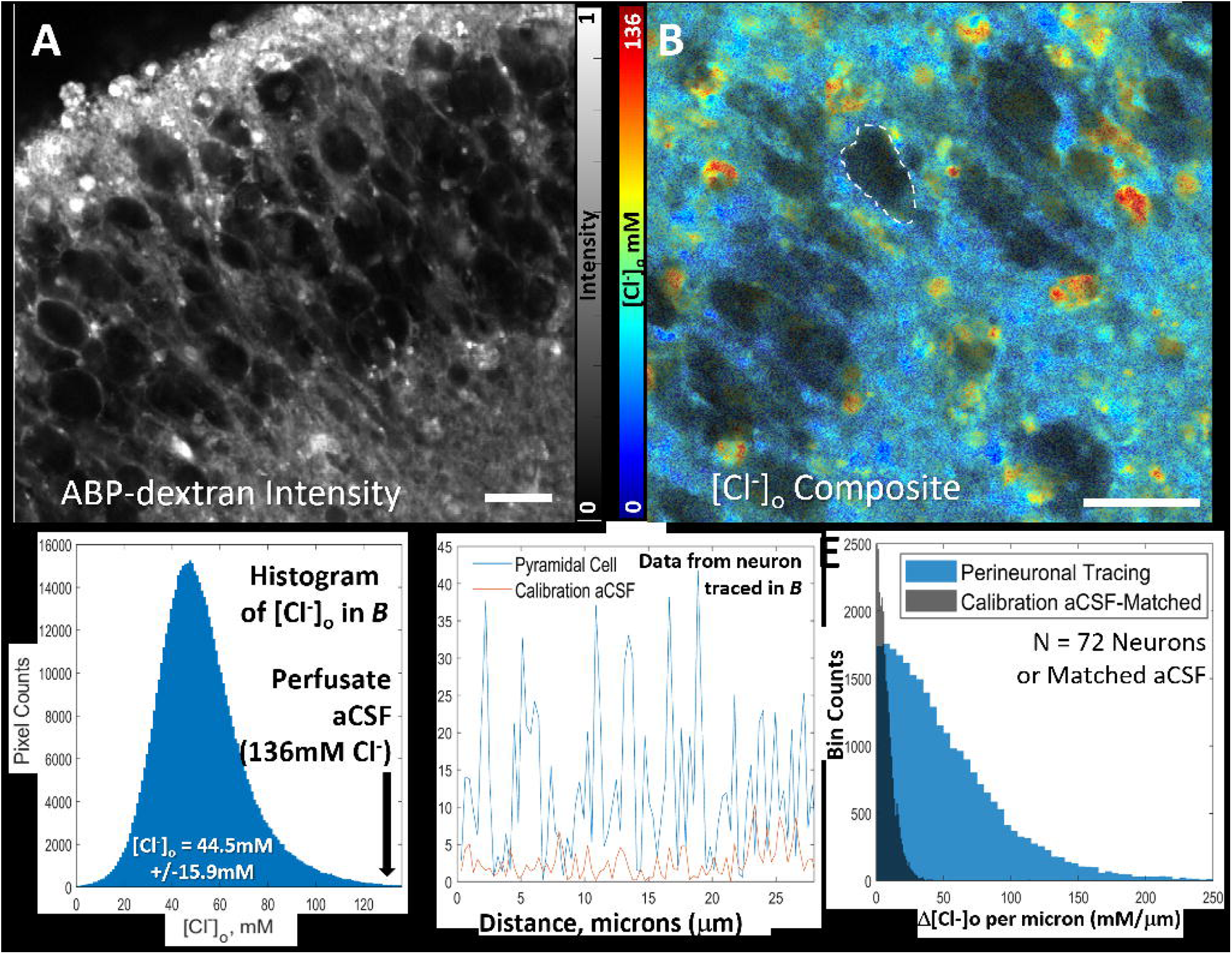
*In vivo* Cl^-^_o_ in mouse neocortex is consistent with *ex vivo* findings. ***A&B)*** Schematic of *in vivo* imaging set-up, using a surgically implanted headbar to immobilize P26- P30 mouse pup for imaging through 3mm coverglass window overlaying 1% agar brought up in aCSF containing ABP-Dextran and 136mM chloride laid directly on exposed cortex. ***C)*** Intensity image of mouse L1 cortex (depth 54μm) demonstrating ABP-dextran diffusion into parenchyma and exclusion from vessels. ***D)*** Intensity image of L2/3 cortex 138μm deep to ***C***, demonstrating ABP-dextran exclusion from cells and vessels; large vessels appear blurred because their widest diameter exists superficial to this field (depth 192μm). Inset is 7x zoomed portion of corresponding chloride composite image, one of three fields from this animal from which we identified and analyzed pyramidal neurons in a manner similar to Figure 3. ***E)*** Region of interest (ROI) selections shown numbered in Cl^-^_o_ composite image, corresponding to intensity image in ***C***, demonstrates exclusion of large vessels from ROIs but is otherwise permissive. ***F)*** Change in chloride (Δ[Cl^-^]_o_) plotted against distance in the perineuronal space (blue line) or the same trace overlaid onto a calibration aCSF FLIM image (red line) of the representative neuron traced in ***D*** inset (white dashes). ***G)*** Perineuronal Δ[Cl^-^]_o_ per micron calculated from 11 L2/3 neurons’ circumferential traces demonstrates that the variability of perineuronal Cl^-^_o_ *in vivo* exceeds the variability of control aCSF (p < 0.001, by Kolmogorov-Smirnov test; *N* > 1,030 pixel-wise measurements from 11 neurons from 3 fields of 1 animal; see *Methods*), consistent with our *in vitro* findings. ***H)*** Cl^-^_o_ in mouse neocortex is significantly lower than the CSF Cl^-^ as shown in this violin plot of all ROI pixels with population mean (black dashed lines) and overlaid mean+/- standard deviation of ROIs (blue circles with red bars) analyzed. Compared to agarose brought up in 136mM aCSF overlying the cortex (leftmost violin plot: 135.4+/-5.0mM; mean+/-standard deviation of all analyzed pixels; 135.4+/-0.8mM, mean+/-standard deviation of *N* = 4 ROIs), *in vivo* mouse cortex had significantly lower Cl^-^_o_ whether measured in cellular fields such as in ***C*** or in acellular fields such as in ***D*** (plots 2 and 3: 48.9+/-6.7mM and 55.0+/-17.9mM pixel-wise, respectively; 47.2+/-2.4mM of *N* = 5 ROIs and 49.2+/-2.7mM of *N* = 7 ROIs, respectively). This was true when considering all ROIs in all fields from the animal from which ***C&D*** were taken (plot 4: 47.2+/-14.9mM pixel-wise; 46.2+/-8.0mM of *N* = 28 ROIs), and when considering all fields from all animals used in this study (plot 5: 48.1+/-17.0mM; 49.4+/-11.2mM of *N* = 68 ROIs). Statistically comparing discreet ROI means by ANOVA analysis with Bonferroni correction, all groups demonstrated a statistically significant difference vs control (p < 0.001) but were not significantly different vs each other (p = 1.0000). Note that ROIs are too numerous to display for all groups, and instead the mean+/-standard deviation (blue circles +/- red lines) calculated from ROI mean values is overlaid onto the violin plots. Violin plot means (dashed black lines), reflecting the mean of aggregated pixel-wise Cl^-^_o_ values from all ROIs, appear on the graph; in the interest of clarity, associated standard deviations are listed above but are not explicitly labeled on the graph. *Scale bars **C,D&E**: 50μm; inset scale bar D: 5μm; significance values noted graphically for p-values <0.05 (*); <0.01 (**); <0.001 (***); “ns” indicates “not significant”*.

## Discussion

We demonstrate stable Donnan exclusion of chloride by fixed anions in a transport-free system of sulfated agar using FLIM of a novel fluorophore, ABP-dextran, and validate these measurements with direct chemical assays of sulfate and chloride (Figure 2). FLIM of ABP- dextran demonstrated that Cl^-^_o_ in the brain is less than half the Cl concentration in the cerebrospinal fluid and plasma (Figures 3-6). This low Cl^-^_o_ arises from Donnan exclusion of chloride by fixed anions including sulfated glycosaminoglycans (sGAGs), because release of sGAGs by chondroitinase results in an increase in Cl^-^_o_ that varies with the amount of sulfates released (Figure 5). Major findings are confirmed *in vivo* (Figure 6).

### Limitations of this study

The finding that Cl^-^_o_ is much lower than expected in the extracellular space of living brain tissue is based on FLIM of a new Cl-sensitive fluorophore, ABP-dextran (Normoyle *et al*., 2024). These results are in contrast to the higher values measured using ion sensitive electrodes (Jiang *et al*., 1992; Kroeger *et al*., 2010), a discrepancy that we cannot definitively explain at this time. The ABP-dextran fluorophore is based on a validated structure (Biwersi *et al*., 1992) and was calibrated against Cl, other anions, and direct chemical analysis of chloride concentrations *in situ* (Figures 1&2). ABP-dextran FLIM accurately reported Cl values in sulfated carbohydrate polymers (Figure 2), so it is unlikely that the gangliosides, sGAGs or other anions in the extracellular environment (Figure 1C) affected the accuracy of the ABP-dextran FLIM. ABP- dextran overcomes several barriers to accurate measurement of extracellular chloride, including sufficient sensitivity in the relevant range of chloride concentrations, competition with autofluorescence, and exclusion from the cytoplasmic space, where lower chloride values would be encountered. ABP-dextran was clearly excluded from the intracellular space (Figure 3A&B; Figure 4A&C; Figure 6D) and blood vessels (Figure 6C,D&E). Experiments were focused on establishing the utility of ABP-dextran as an extracellular-specific sensor and did not include transgenic fluorophores that could have measured Cl^-^_i_ and marked neurons for *in vivo* analysis. We speculate that the difference between our non-invasive, novel fluorophore-based measures and the implanted ion-sensitive electrode measures of Cl^-^_o_ may arise from the large difference in sample sizes, where microelectrodes sample a single microenvironment while the fluorometric technique samples millions. It is much easier to perceive the wide distribution of Cl^-^_o_ with the larger sample size (Figure 4D), and to appreciate the average Cl^-^_o_. A study directly comparing fluorometric and microelectrode techniques would be of great interest.

### Stable Donnan exclusion

An important finding in this study is that local Cl differences established by Donnan exclusion are stable for weeks, and do not require active transport to be maintained (Figure 2). This phenomenon is well-established in biophysics (Epsztein *et al*., 2018; Gao *et al*., 2022), but is more confusing in neuronal physiology where membranes with time-varying ionic permeabilities and transporters create a more complex environment (Voipio *et al*., 2014; Doyon *et al*., 2016; Düsterwald *et al*., 2018; Rahmati *et al*., 2021). A point of particular confusion is the role of active Cl transport vs. Donnan exclusion in maintaining a particular E_GABA_ in the face of GABA-gated Cl flux that alters the local Cl concentration. After a GABA-gated Cl influx, and re-establishment of local charge balance by cationic flux via voltage- and ligand-gated cation channels, the local cytoplasm will contain excess chloride salts. This Cl has two fates: either excess salt diffuses through the cytoplasm to re-establish a new equilibrium with fixed charges, or the excess salt is transported back across the membrane by cation-chloride cotransporters (Brumback & Staley, 2008; Lewin *et al*., 2012; Doyon *et al*., 2016; Beckstein & Naughton, 2022). The transporters are required to re-establish the original steady state volume, but in the absence of transport, diffusion maintains local Cl concentrations remarkably well (Brumback & Staley, 2008; Rahmati *et al*., 2021). In the absence of membrane Cl salt transport, this diffusion of excess Cl salts will be associated with either a cytoplasmic volume increase from associated water influx or an increase in the mean Cl concentration in the cytoplasm. At pathologically high rates of Cl influx, membrane chloride salt transport becomes rate-limiting (Staley & Proctor, 1999; Jedlicka *et al*., 2011; Lewin *et al*., 2012; Doyon *et al*., 2016), and chloride concentrations become sufficiently labile to depolarize E_GABA_ (Barker & Ransom, 1978; Alger & Nicoll, 1982; Huguenard & Alger, 1986; Staley *et al*., 1995). Thus Donnan exclusion *establishes* the local value of E_GABA_, and active transport *maintains* that value (and neuronal volume) in the face of synaptic Cl flux (Glykys *et al*., 2014a). The same principals should apply to the local extracellular chloride concentrations described here, with astrocytic chloride buffering (Egawa *et al*., 2013) and diffusion between the extracellular fluid, the CSF, and vascular spaces (Rasmussen *et al*., 2022) taking the place of neuronal membrane cation-chloride cotransport. Importantly, because E_GABA_ is set by the ratio of Cl^-^_o_ to Cl^-^_i_ (as given by the Nernst equation) and neurons can actively adjust intracellular immobile anions (Rahmati *et al*., 2021), there is no Cl^-^_o_ level that expressly precludes hyperpolarizing inhibition because proportional Cl^-^_i_ reduction allows E_GABA_ to be maintained at any given value (Figure 7). Multi-ion modeling from our group has previously demonstrated similarly modest changes in E_GABA_ with Cl^-^_o_ reduction that can be counteracted by even more modest Cl^-^_i_ changes (Glykys *et al*., 2017).

**Figure 7.**
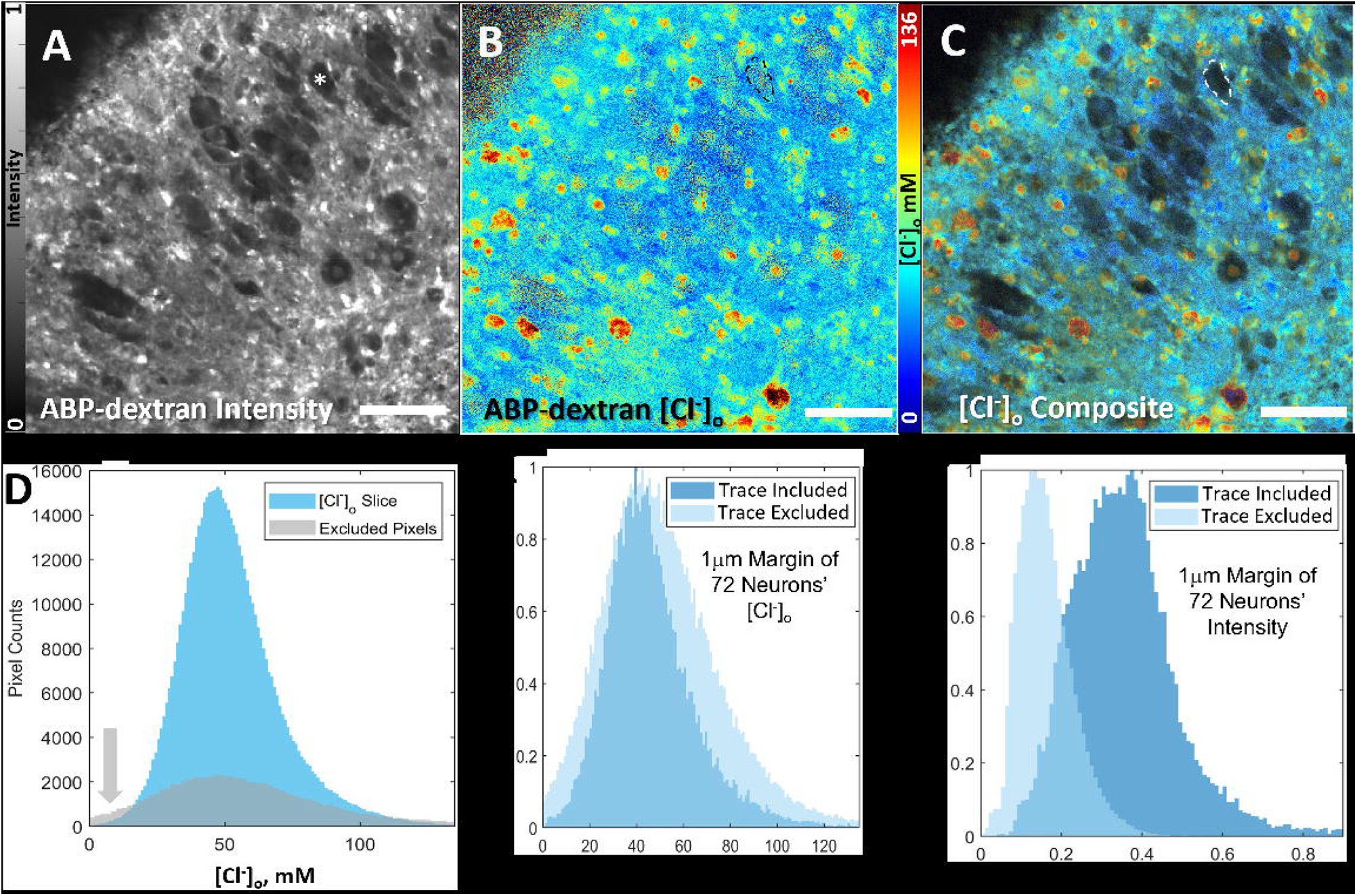
E_GABA_ is maintained at any Cl^-^ given proportionate change in Cl^-^. The reversal potential of chloride across GABA_A_Rs (E_GABA_) is given by the Nernst equation, 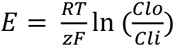 At constant temperature the only variables that determine E_GABA_ are Cl^-^ and Cl^-^, or rather the only single variable is the ratio of Cl^-^_o_ to Cl^-^_i_. Using the Nernst equation to model E_GABA_ across the physiological range of Cl^-^_i_ at several Cl^-^_o_ values, we unsurprisingly find that hyperpolarizing E_GABA_ values can be achieved at any Cl^-^_o_ value. Lower Cl^-^_o_ values simply need a proportionately lower Cl^-^_i_ value to achieve a given E_GABA_, which neurons are capable of by increasing intracellular immobile anions such as cytoskeletal biopolymers (Rahmati *et al*., 2021). In the examples given above, the Cl^-^_i_ necessary to maintain E_GABA_ of −80mV (dashed red line) varies from 1mM to 6mM as Cl^-^_o_ is varied from 20mM to 120mM. More complex multi-ion modeling will be facilitated with higher resolution and spatiotemporally correlated measurements of Cl^-^_o_ and Cl^-^_i_ in proximity across the neuronal membrane.

### Low extracellular Cl

The Cl^-^_o_ measured by ABP FLIM is less than half of the canonical Cl^-^_o_. Such a low value of Cl^-^_o_ should affect not only E_GABA_ but also the inhibitory GABA_A_ conductance, because there are only half as many ions to carry the outward membrane currents (i.e. inward Cl- flux) (Coombs *et al*., 1955). Shouldn’t such a low value of Cl^-^_o_ have been obvious from electrophysiological studies? E_GABA_ is determined by the ratio of Cl^-^_i_ to Cl^-^_o_ (Coombs *et al*., 1955; Staley, 2023). If either concentration is known, the other can be determined from E_GABA_. However if both Cl^-^_i_ and Cl^-^_o_ are determined by the concentrations of local immobile anions, neither can be set experimentally and thus E_GABA_ cannot be used to determine either Cl^-^_i_ or Cl^-^_o_; only the ratio can be inferred from the Nernst equation. Similarly, although the conductance of inward Cl flux (outward membrane current) is determined in part by Cl^-^_o,_ it is also determined by the GABA_A_ permeability, i.e. the number of open membrane channels and their capacity for ion flux. Thus membrane currents are not sufficient to determine Cl^-^_o_ (Coombs *et al*., 1955). Control of the ionic milieu is greater in single-channel recordings, but whether the complement of immobile anions is physiological under these conditions is difficult to determine, and even in these conditions there is substantial uncertainty regarding local ionic concentrations (Li *et al*., 2015). The effects of variable Cl^-^_i_ and Cl^-^_o_ have on E_GABA_ have been explored (Glykys *et al*., 2017), but high resolution spatiotemporal correlation of all three parameters will provide exciting new insights into dendritic signal processing.

### Cl^-^_o_ displacement

The extracellular matrix of the brain is often depicted as perineural nets surrounding parvalbumin neurons that are detected by specific agglutinins and antibodies (Celio & Blümcke, 1994). However, these assays detect very specific motifs of glycoproteins and glycosaminoglycan sulfation; using more permissive ionic methods, a highly anionic extracellular matrix can be seen to surround virtually all neurons (Golgi, 1898; Glykys *et al*., 2014a; Morawski *et al*., 2015). This extracellular matrix fills the extracellular space, which comprises about 20% of the brain’s volume (Syková & Nicholson, 2008; Tønnesen *et al*., 2018). We have focused our analysis on the sGAG moiety of proteoglycans because these are prominent constituents of perineuronal extracellular matrix (Tønnesen *et al*., 2018; Karamanos *et al*., 2021), are found in both dense and more diffuse neuronal extracellular matrix (Deepa *et al*., 2006), and are readily perturbed experimentally with commercially available enzymes. GAGs are polymers of widely varied length comprised of sugars and substituted sugars (N-acetylglucosamine, glucuronic acid, and galactose) that are synthesized without templates. Up to 3 sulfates are added to the sugar moieties before export to the extracellular space. This enables the creation of a stable but highly variable spatial distribution of dense, fixed anionic charge in the extracellular matrix. Figure 3 demonstrates that this variance is substantial at the level of resolution of light microscopy. We have previously demonstrated reduction of fixed extracellular anions after chondroitinase treatment (Glykys et al., 2014a), and others have demonstrated small but significant depolarizing shifts of E_GABA_ after chondroitinase treatment (Klein et al., 2018). However, the distribution of fixed anions at the extracellular face of GABA_A_ receptors, and the corresponding Cl^-^_o_, is currently unknown. The effects of variable Cl^-^_o_ and Cl^-^_i_ have on E_GABA_ have been explored (Glykys *et al*., 2017), but high resolution spatiotemporal correlation of all three variables would allow more descriptive modeling in complex solutions.

### Physiological implications

Along with Cl^-^_i_ microdomains (Rahmati *et al*., 2021), Cl^-^_o_ microdomains in the region of the extracellular openings of GABA_A_ receptor-operated channels could be a critical determinant of both E_GABA_ and the GABA_A_ conductance. The wide range of Cl^-^_o_ values established via Donnan exclusion by sulfated GAGs (Figures 2, 3, 4, 5 and 6) makes possible a substantial variety of effects of GABA_A_ receptor activation based on the direction of chloride flux. These effects range from hyperpolarizing inhibition through shunting inhibition through excitation by activation of low threshold calcium currents and removal of the magnesium block of the NMDA receptor. In addition, the effects of Cl^-^_o_ on the quantity of charge that is carried by GABA_A_ membrane currents (the conductance) (Coombs *et al*., 1955) makes possible a broad range of strengths of GABA_A_ synapses that is independent of the number or subunit composition of receptors or their posttranslational modifications.

The turnover of the macromolecules comprising the extracellular matrix is very slow (Tsien, 2013), on the order of months to years based on rodent experiments (Margolis & Margolis, 1973) as well as the rate of deterioration of patients with genetic defects of GAG catabolism (Héron *et al*., 2011; Hampe *et al*., 2020; Seo *et al*., 2020). This slow turnover provides a mechanism for long-lasting influences on synaptic GABA signaling. Some elements of GAG synthesis are activity dependent (Sidharthan *et al*., 2013), but currently nothing is known about the insertion of GAGs into the extracellular matrix, in situ modification, and the capacity for directed alterations of Cl^-^_o_ in the regions of GABA_A_ receptors and synapses. Thus the role of Cl^-^_o_ in synaptic signal processing and memory remains to be discovered.

### Pathological implications

Cytotoxic cerebral edema: If Cl^-^_o_ is normally only half of the value of Cl in CSF and plasma, then the capacity for Cl^-^_o_ to increase acutely may be an important and hitherto unappreciated element of the pathophysiology of cerebral edema (Glykys *et al*., 2017). Matrix metalloproteinases (MMPs) are released and activated after neuronal injury and death (Zhang *et al*., 2016). Release of sulfates in sGAGs from the extracellular matrix would reduce the Donnan exclusion of Cl by fixed sulfates. This would result in an acute increase of Cl^-^_o_ as in Figure 4. The action of equilibrative cation-Cl cotransporters would then lead to a corresponding increase in chloride salts in the cytoplasm of neighboring healthy neurons, with attendant volume increases. This novel mechanism of cytotoxic edema could potentially be ameliorated by pharmacological MMP inhibition. Alternatively, inhibition of the influx of chloride salts that cross the BBB from the vascular space (Jha *et al*., 2019) would limit the influx of Cl into the extra and intracellular spaces.

Post Traumatic Epilepsy: Richard Miles and colleagues demonstrated that GABA_A_ receptor activation could be excitatory in human brain tissue resected for control of medically intractable epilepsy (Cohen *et al*., 2002). However, the pathomechanism has never been elucidated (Huberfeld *et al*., 2007; González, 2016; Karlócai *et al*., 2016). Gliosis is the most common finding in medically intractable epilepsy (Thom, 2014). In glial scars, reactive astrocytes synthesize new matrix elements (Roll & Faissner, 2014; Song & Dityatev, 2018). If the replacement matrix was more heavily sulfated than the original, as would be expected from the firm texture that is responsible for the name “scar” (Lesperance *et al*., 1992), then in areas of gliosis Cl^-^_o_ would be low. Low Cl^-^_o_ could contribute to degradation of GABA_A_ receptor mediated inhibition, frank excitation, and failure of anticonvulsants whose action are predicated on intact synaptic inhibition. Thus replacement of the extracellular matrix and distortion of Cl^-^_o_ could comprise an important mechanism of medically intractable epilepsy after brain injury.

### Future studies

The findings that Cl^-^_o_ is both lower than expected and spatially heterogenous opens the door for many investigations into the mechanisms by which Cl^-^_o_ is set, such as the synthesis, export, and in situ modification of sulfated GAGs. The activity dependence of these mechanisms and the effect of Cl^-^_o_ on synaptic signaling are additional areas of investigation. Pathologically, the impact of changes in Cl^-^_o_ in both acute edema and long-term complications such as epilepsy may provide much-needed therapeutic insights into these frequently-intractable conditions.

## Abstract Figure Legend

The perineuronal space contains high concentrations of immobile glycoproteins and glycolipids bearing sialic acid (such as glycosides) and sulfate groups (such as sulfatides and glycosaminoglycans) that together account for more than 50 mEq/L anionic charge. ***Left side***: Ion concentrations in the perineuronal space are typically treated as equivalent to cerebrospinal fluid (CSF) values, requiring that the component ion concentrations of CSF are unaltered in the extracellular space. ***Right side***: However, the large pool of membrane-associated immobile anionic species that account for nearly half the total extracellular anionic charge results in Donnan exclusion, redistributing the mobile anionic charge to maintain a constant total charge (immobile plus mobile charges) in accordance with Gauss’ Law of electroneutrality. Note that the same number of positive charges are present, but that the presence of the rectangular structures representing perineuronal membrane-associated immobile anions – from sialic acid-bearing glycosides to chondroitin sulfate-decorated aggrecan family proteoglycans – make up roughly half the total anionic charge. The total extracellular charge is the same on the left and right sides of the schematic, both for cations and anions, even though the number of mobile anions on the right is half the number on the left. Because the immobile anions are known to be heterogeneously distributed, Gauss’ Law implies that mobile anions, principally chloride, must also be heterogeneously distributed in the extracellular space to maintain total anionic charge. Because transmembrane chloride flux underlies GABAergic inhibition, we directly and non-invasively measure extracellular chloride to better understand the influence of Donnan exclusion on chloride distribution and inhibitory function *in vitro* and *in vivo*.

## Additional Information

### Competing Interests

None of the authors have any competing interests to declare, as indicated during the submission process for each author individually.

### Funding

This work was funded by three grants to Kevin J. Staley from the National Institute for Neurological Disorders and Stroke (NINDS): NIH R01 NS040109, R35 NS116852 and P01 NS127769.

### Author Contributions

KPN, KPL, KE and KJS conceived of and designed the approach and execution of these experiments. KPN and KJS are responsible for specific experimental design, interpretation of experiments, and manuscript preparation. KPL designed and built the two-photon microscopes and operational software that made this work possible. MAM and LL lent expertise and support for in vivo experiments, including training KPN in mouse cranial window surgery. LL and FHS provided expertise and critical input during experimental interpretation and manuscript writing. MP and BPC designed and executed HPLC experiments specific to this manuscript. This research was made possible by NINDS funding to KJS (NIH R01 NS040109, R35 NS116852 and P01 NS127769).

### Authors’ Translational Perspective

We hypothesized that if measured non-invasively, the extracellular chloride concentration (Cl^-^_o_) would be less than CSF chloride values (≈110mM) due to Donnan exclusion by heterogeneously distributed extracellular anions. We found that Cl^-^_o_ has a mean value much lower than previously appreciated (≈50mM). Our data further demonstrate that the loss of sulfated glycosaminoglycans (sGAGs) from the extracellular matrix results in an increased Cl^-^_o_ due to decreased Donnan exclusion. Together with our previous work demonstrating heterogenous intracellular chloride (Cl^-^_i_), our data suggests the variance of GABA reversal potential (E_GABA_) we and others have observed is due to the heterogeneity of the concentrations of the primary permeant anion, Cl^-^, on both the intra- and extracellular side of the channel. Stable variations of perisynaptic Cl^-^_o_ or Cl^-^_i_ potentially allows for the tuning of E_GABA_ at individual GABA_A_ synapses and suggests a mechanism for information storage.

Pathophysiological alteration of Cl^-^_o_ would be expected to occur when the extracellular matrix is damaged, such as when matrix metalloproteases (MMPs) are released following neuronal injury or death. The expected increase in extracellular chloride salts, in turn leading to equilibrative intracellular increases, represents a new mechanism of cytotoxic edema.

## Acknowledgements

This work was supported by NIH R01 NS040109, NIH R35 NS116852, and NIH P01 NS127769 to KJS. The authors would like to thank current and former members of the Pediatric Epilepsy Research Laboratory at MGH for critical input, discussions, and support during experimental development including Joseph Glykys, Negah Rahmati, Trevor Balena, Rehan Raiyyani, and Michelle Mail. The authors would also like to thank members of the MGH Institute for Neurodegenerative Disorders, particularly Steven S. Hou, for technical expertise and related advice.

## Notes

https://github.com/KieranNormoyle/NeuronalChloride2023

### Competing Interest Statement

The authors have declared no competing interest.

### Summary of Updates

This revision reflects inclusion of modeling data (Figure 7) and new analyses stemming from a final round of review.

